# The physical chemistry of interphase loop extrusion

**DOI:** 10.1101/2024.08.23.609419

**Authors:** Maxime M.C. Tortora, Geoffrey Fudenberg

## Abstract

Loop extrusion constitutes a universal mechanism of genome organization, whereby structural maintenance of chromosomes (SMC) protein complexes load onto the chromatin fiber and generate DNA loops of increasingly-larger sizes until their eventual release. In mammalian interphase cells, loop extrusion is mediated by the cohesin complex, which is dynamically regulated by the interchange of multiple accessory proteins. Although these regulators bind the core cohesin complex only transiently, their disruption can dramatically alter cohesin dynamics, gene expression, chromosome morphology and contact patterns. Still, a theory of how cohesin regulators and their molecular interplay with the core complex modulate genome folding remains at large. Here we derive a model of cohesin loop extrusion from first principles, based on *in vivo* measurements of the abundance and dynamics of cohesin regulators. We systematically evaluate potential chemical reaction networks that describe the association of cohesin with its regulators and with the chromatin fiber. Remarkably, experimental observations are consistent with only a single biochemical reaction cycle, which results in a unique minimal model that may be fully parameterized by quantitative protein measurements. We demonstrate how distinct roles for cohesin regulators emerge simply from the structure of the reaction network, and how their dynamic exchange can regulate loop extrusion kinetics over time-scales that far exceed their own chromatin residence times. By embedding our cohesin biochemical reaction network within biophysical chromatin simulations, we evidence how variations in regulatory protein abundance can alter chromatin architecture across multiple length- and time-scales. Predictions from our model are corroborated by biophysical and biochemical assays, optical microscopy observations, and Hi-C conformation capture techniques. More broadly, our theoretical and numerical framework bridges the gap between *in vitro* observations of extrusion motor dynamics at the molecular scale and their structural consequences at the genome-wide level.

## Introduction

Genomes are continuously and actively organized by loop extrusion (Dekker et al., 2023). During this stochastic process, molecular motors land on DNA and generate progressively enlarging loops, until they eventually dissociate (Alipour and Marko, 2012). Strong support for loop extrusion *in vivo* stems from the comparison of polymer model predictions with genomics data obtained from Hi-C experiments (Fudenberg et al., 2017), as well as overall chromosome morphology in mitosis (Gibcus et al., 2018; Goloborodko et al., 2016). More recently, *in vitro* single-molecule tracking assays provided direct evidence that the Structural Maintenance of Chromosome complexes (SMCs) cohesin and condensin can processively generate loops on tethered DNA molecules (Davidson et al., 2019; Golfier et al., 2020; Kim et al., 2019). Active modulation of loop extrusion dynamics is now believed to regulate a growing number of cellular decision mechanisms. These range from controlling promoter choice at the protocadherin locus to modulate neural wiring (Kiefer et al., 2023), to VDJ recombination to enable immune repertoire diversity (Hill et al., 2020).

In mammalian interphase cells, cohesin acts as the main loop extruder (Davidson and Peters, 2021). Increasing evidence argues that cohesin cannot be thought of as a monolithic complex, and that transient associations with regulatory proteins modulate the dynamics of loop extrusion (Davidson and Peters, 2021; Pezic et al., 2017). The core cohesin complex consists of SMC1 and SMC3, one of the HEAT-repeat subunits SA1 and SA2, along with the kleisin subunit RAD21. Among these core components, RAD21 acts as the “nexus” (Muir et al., 2016) or “docking point” (Huis in ’t Veld et al., 2014) for the recruitment of cohesin regulatory factors, which is mediated by the presence of multiple interfaces competent for the binding of cohesin regulators. While many of these regulators were originally identified for their functions in sister chromatid cohesion (Nasmyth, 2011; Onn et al., 2008), their roles for extrusion are now increasingly appreciated (Davidson and Peters, 2021).

Individual disruptions to the cohesin regulators NIPBL, PDS5 and WAPL can induce dramatic changes in cohesin properties and genome organization. The depletion of WAPL (i.e., ΔWAPL) leads to a considerable increase in cohesin residence times and loaded fraction, and results in the lengthwise compaction of entire chromosomes into ‘vermicelli’ that are highly enriched for cohesin along their axes (Tedeschi et al., 2013). These chromatids have a prophase-like appearance, yet emerge from the action of interphase cohesin complexes. Similar phenotypes have been reported upon depletion of PDS5 (van Ruiten et al., 2022; Wutz et al., 2017) and overexpression of RAD21 (Sun et al., 2023), but are inhibited by the removal of NIPBL (Haarhuis et al., 2017).

At the molecular level, dissecting the respective functions of cohesin regulators poses a substantial challenge, due to the multiple roles reported for individual regulators. NIPBL has been reported to act as a cohesin loader (Ladurner et al., 2016), but is also required for ATP hydrolysis and translocation *in vitro* (Davidson et al., 2019; Kim et al., 2019). PDS5 may facilitate cohesin unloading in conjunction with WAPL (Ouyang et al. 2013), but also competes with NIPBL by binding with mutual exclusivity to the cohesin complex (Kikuchi et al., 2016; Petela et al., 2018). Existing models describing the function of individual regulatory proteins are largely qualitative, and do not lend themselves to quantitative predictions of their genome-wide consequences *in vivo*. For instance, although PDS5 has been recently identified as a “brake” for loop extrusion (van Ruiten et al., 2022), a mathematical description of how the cohesin extrusion rate depends on the abundance of PDS5, or any other regulator, is currently lacking.

Despite their striking effects on loop extrusion dynamics, biophysical measurements indicate that the residence time of NIPBL, WAPL, and PDS5 on chromatin (∼1min, (Ladurner et al., 2016; Rhodes et al., 2017)) is considerably less than that of cohesin (∼20mins for RAD21, (Gerlich et al., 2006; Hansen et al., 2017; Tedeschi et al., 2013)). This dynamic turnover of regulators on the core cohesin complex implies that quantitative models of loop extrusion would benefit from depicting cohesins as multi-state motors with heterogeneous properties arising from the binding of distinct regulators (Corsi et al., 2023). However, current extrusion models: (i) lack a molecular basis for the roles of different cohesin regulators, and (ii) assume that all cohesins are single-state motors with identical extrusion behavior (Brackey et al., 2020). Because of this, loop extrusion parameters for existing models are obtained in part by fitting to match Hi-C data, rather than coming from first principles or biophysical measurements. Ultimately, new models are needed to incorporate insights from *in vitro* motor assays, account for roles of different cohesin regulators, and understand their downstream impacts on chromosome organization and genomic functions.

Here, we introduce a five-state biochemical reaction network model describing interphase loop extrusion by cohesin. Using available biophysical data from unperturbed cells, this model can predict both the changes in extrusion dynamics observed after depletions of cohesin regulators — as well as their consequences for 3D genome folding — directly from their abundances, without requiring any Hi-C data as input. Our model yields molecular insights into the differential roles of NIPBL, PDS5 and WAPL, and provides a general framework for encoding biophysical data on cohesin and its regulators into computational models of loop extrusion.

## Results

### Building a minimal biochemical reaction network for interphase extrusion

We set out to develop a minimal model of interphase cohesin biochemistry based on mass action kinetics that accounted for the individual roles of the cohesin regulators NIPBL, PDS5, and WAPL. Building on experimental observations (**Methods**), we made three main simplifying assumptions: (i) RAD21 loading and unloading dynamics can be taken as a proxy for those of the core cohesin complex; (ii) regulatory proteins require the core complex to be chromatin-associated; and (iii) regulators bind mutually exclusively to the core complex. Based on their reported absolute nuclear abundances, the number of loaded cohesins exceeds the combined populations of bound NIPBL, PDS5 and WAPL in HeLa cells, implying that cohesin likely exists in a loaded state without regulators, which indicates that a minimal network should include four distinct loaded states and an unloaded state. Note that while some of these approximations are likely too strong — e.g., they do not explicitly consider the possibility of a PDS5-WAPL complex on cohesin (Gandhi et al., 2006; Kueng et al., 2006) — they yield a tractable description of the cohesin regulatory network that we demonstrate is sufficient to recapitulate a broad range of experimental observations.

To determine reaction network transition rates, we curated experimental measurements of protein abundance, chromatin bound fraction, and residence times for RAD21, NIPBL, WAPL, and PDS5 in HeLa cells (amalgamating paralogs PDS5A and PDS5B, **Table 1; Methods**). The bound fraction and residence times for each of the 4 proteins from Fluorescence Recovery After Photobleaching (FRAP) enabled us to define and uniquely determine a rich zoo of minimal reaction networks with up to 8 chemical transition rates (**Supp. Sec. I**). The simplest class of such reaction networks are completely reversible and acyclic, and are characterized by linear, branched, and star topologies (**Fig. S1**). However, none of these networks provided viable descriptions of cohesin biochemistry, as none of them predict the experimentally observed increase in the bound fraction of RAD21 upon WAPL depletion (Haarhuis et al., 2017; Tedeschi et al., 2013; Wutz et al., 2017). We thus provide mathematical evidence that the chromatin entry and exit of cohesin via distinct molecular pathways applies not only for sister chromatid cohesion in S phase (Nasmyth, 2011; Onn et al., 2008), but also holds for loop extrusion in interphase.

**Table 1:** Absolute nuclear copy numbers, chromatin bound fractions and residence times of each protein used to constrain the model in wild-type HeLa cells (see Methods).

| Protein | Copy number | Chromatin bound % | Chromatin residence time |
| --- | --- | --- | --- |
| RAD21 | 264,000<br>(Bekker-Jensen et al., 2017; Holzmann et al., 2019) | 65 %<br>(Holzmann et al. 2019) | 822 s<br>(Holzmann et al. 2019) |
| NIPBL | 111,000<br>(Bekker-Jensen et al., 2017; Holzmann et al., 2019) | 40 %<br>(Rhodes et al., 2017) | 72 s<br>(Rhodes et al., 2017) |
| WAPL | 65,000<br>(Bekker-Jensen et al., 2017; Holzmann et al., 2019) | 35 %<br>(Ladurner et al., 2016) | 45 s<br>(Ladurner et al., 2016) |
| PDS5A/B | 164,000<br>(Bekker-Jensen et al., 2017; Ding et al., 2011) | 45 %<br>(Ladurner et al., 2016) | 70 s<br>(Ladurner et al., 2016) |

We next considered models describing cohesin biochemistry as a reaction cycle with irreversible loading and unloading transitions. For these reaction networks, loading of the core complex onto chromatin occurs via a one-way, irreversible, transition – potentially concomitant with the co-binding of a regulatory protein. After loading, additional cohesin regulators may reversibly bind and unbind the loaded core complex, until its eventual irreversible unloading (**Fig. 1a**). Such cyclic reaction networks confer distinct roles upon regulatory proteins based on their position within the cycle: the first co-binding factor acts as the primary loader, while the last co-binding factor acts as the primary unloader. To systematically consider the potential functions of each regulator, we determined the transition rates for each of 24 possible reaction cycles using biophysical data (**Fig. 1b**). We next pruned networks with unphysical chemical kinetics (i.e., negative transition rates between states) to weed out reaction cycles that are incompatible with the experimental binding kinetics of each regulator. We finally required that networks reproduce three qualitative observations for the bound fraction of RAD21, specifically: (i) a decrease after ΔNIPBL; (ii) an increase after ΔPDS5; and (iii) an increase after ΔWAPL (Wutz et al., 2017). Together, our two-stage pruning procedure uncovered a unique reaction cycle consistent with available data (**Fig. 1c**), which we refer to as the “five-state model”. In this five-state model, NIPBL acts as the primary loader and WAPL as the primary unloader. PDS5 is required to recruit WAPL, but plays no direct role in unloading *per se*, and also competes with NIPBL for association with the “bare” loaded cohesin (*R*) state.

**Figure 1:**
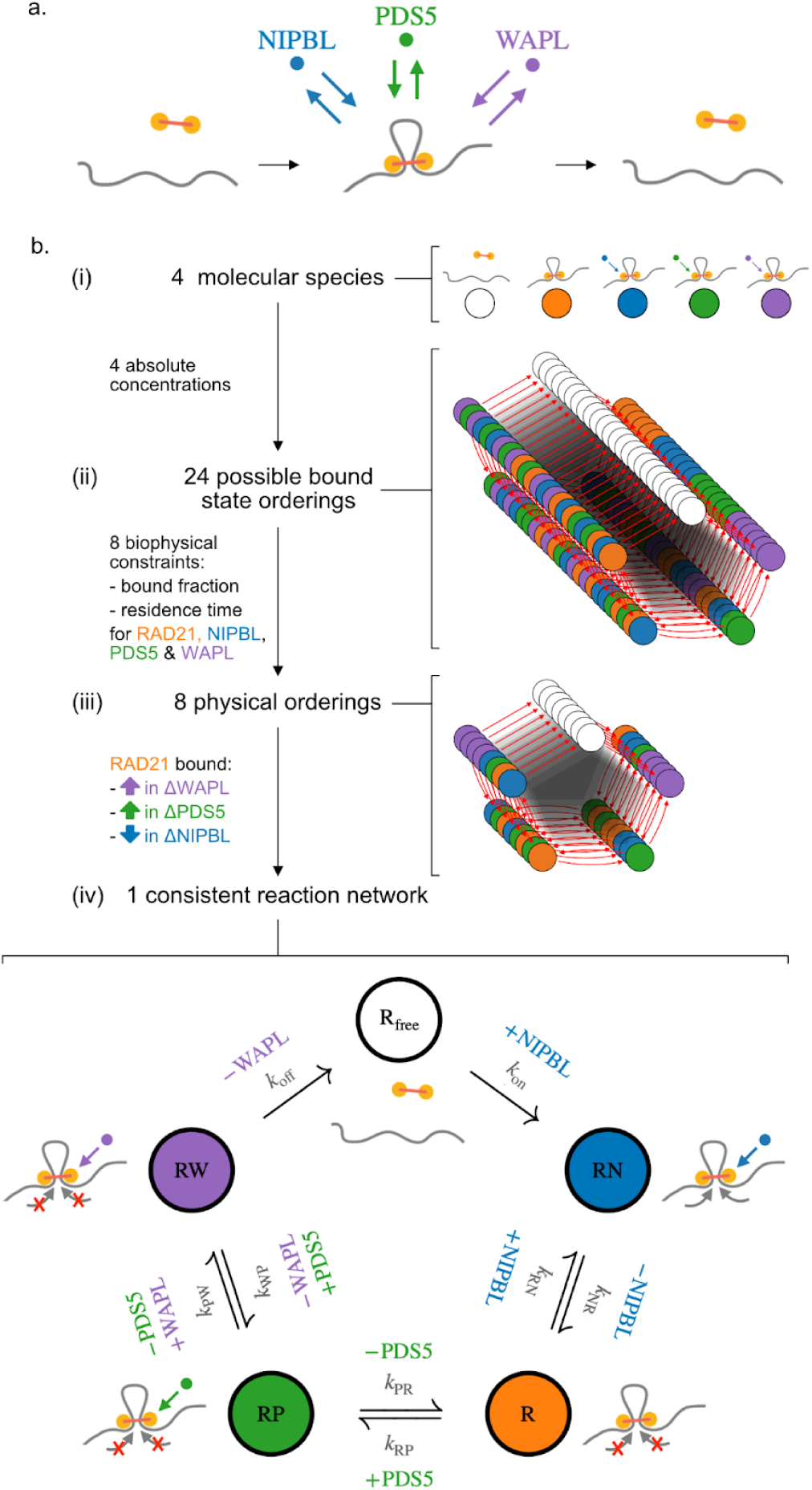
Determining the biochemical reaction network of interphase cohesin. **a.** Illustration of cohesin complex binding, exchange of regulator proteins on the loaded complex at short (∼1 min) timescales, and cohesin unbinding at longer (∼10 min) timescales. **b.** Pruning procedure to obtain a minimal biochemical network model describing the interplay of the regulatory proteins NIPBL, PDS5 & WAPL. ***(i)*** Modeling these three regulators along with the core complex (using RAD21 as its proxy) yields one unloaded and four loaded states. ***(ii)*** Pruning begins by considering all possible sequences (24) of regulator exchange on loaded cohesin complexes. ***(iii)*** After constraining the total nuclear abundance of each protein based on mass spectrometry and FCS measurements in HeLa cells, only 8 reaction networks were physically compatible with the experimental chromatin bound fractions and residence times of RAD21, NIPBL, PDS5 and WAPL as estimated by FRAP in HeLa (see **Methods**). ***(iv)*** Further using the fact that chromatin-associated cohesin increases for ΔWAPL and ΔPDS5, but decreases in ΔNIPBL after RNAi depletion (Wutz et al., 2017) yields a single reaction network (“five-state model”) consistent with *in vivo* observations.

To understand the favorable properties of the five-state model, we tested whether closely-related reaction networks could also be reconciled with experimental data (**Fig. S2**). We first considered a model where, instead of being directly involved in loading, the NIPBL-bound state constitutes an excursion from the main reaction cycle (**Fig. S2a**). For this topology, however, NIPBL depletion does not sufficiently lower the bound fraction of cohesin to agree with experiments (Wutz et al., 2017). The failure of this topology thus supports a direct role for NIBPL in cohesin loading. We next considered a PDS5 excursion model, where PDS5 reversibly binds the core complex but does not recruit WAPL (**Fig. S2b**). With this topology, ΔPDS5 actually slightly lowered RAD21 cohesin residence time, instead of increasing it as observed experimentally (Wutz et al., 2017). The inconsistency of this topology thus argues that PDS5 helps recruit WAPL to promote cohesin unloading.

Together, our minimalistic five-state cohesin reaction network recapitulates biophysical data for RAD21, NIPBL, PDS5 and WAPL, and uniquely identifies their respective roles in the cohesin loading/unloading cycle.

### Coupling the five-state cohesin model with polymer simulations predicts 3D genome folding

To compare predictions of the five-state cohesin model with experimental Hi-C, we used our cohesin reaction network as an input for polymer simulations of chromatin. As for previous approaches (Fudenberg et al., 2016; Nuebler et al., 2018), we coupled a 1D lattice model of cohesin translocation, which tracks cohesin positions over time along the genome, with coarse-grained molecular dynamics (MD) simulations of a generic chromosome, which track chromatin and extruder positions in 3D (**Methods**). In the lattice model, cohesins stochastically load, unload and extrude chromatin. However, we modified cohesin activity in the lattice model to directly depend on the states and transition rates extracted from our five-state model (**Table S1**). To simulate stochastic transitions between states for individual cohesin complexes, we used a discrete-time kinetic Monte-Carlo approach (**Fig. 2a**). Based on the observation that NIPBL is required for ATP hydrolysis by the cohesin complex (Davidson et al., 2019; Kim et al., 2019), we assumed that extrusion occurs only in the NIPBL-bound state (*RN*) (**Fig. 2b**). As hypothesized for condensin-condensin interactions (Samejima et al., 2024), we further assume that cohesin-cohesin encounters along the lattice lead to collisions without bypass. Thus, in the other chromatin-loaded states (*R*, *RP*, *RW*), cohesin remains immobile yet still blocks translocation by active extruders. Extruders are loaded upon transition from the free to NIPBL-bound state and unloaded upon transition from the WAPL-bound to free state. For parsimony, we considered that transition rates between states are homogeneous across all genomic positions.

**Figure 2:**
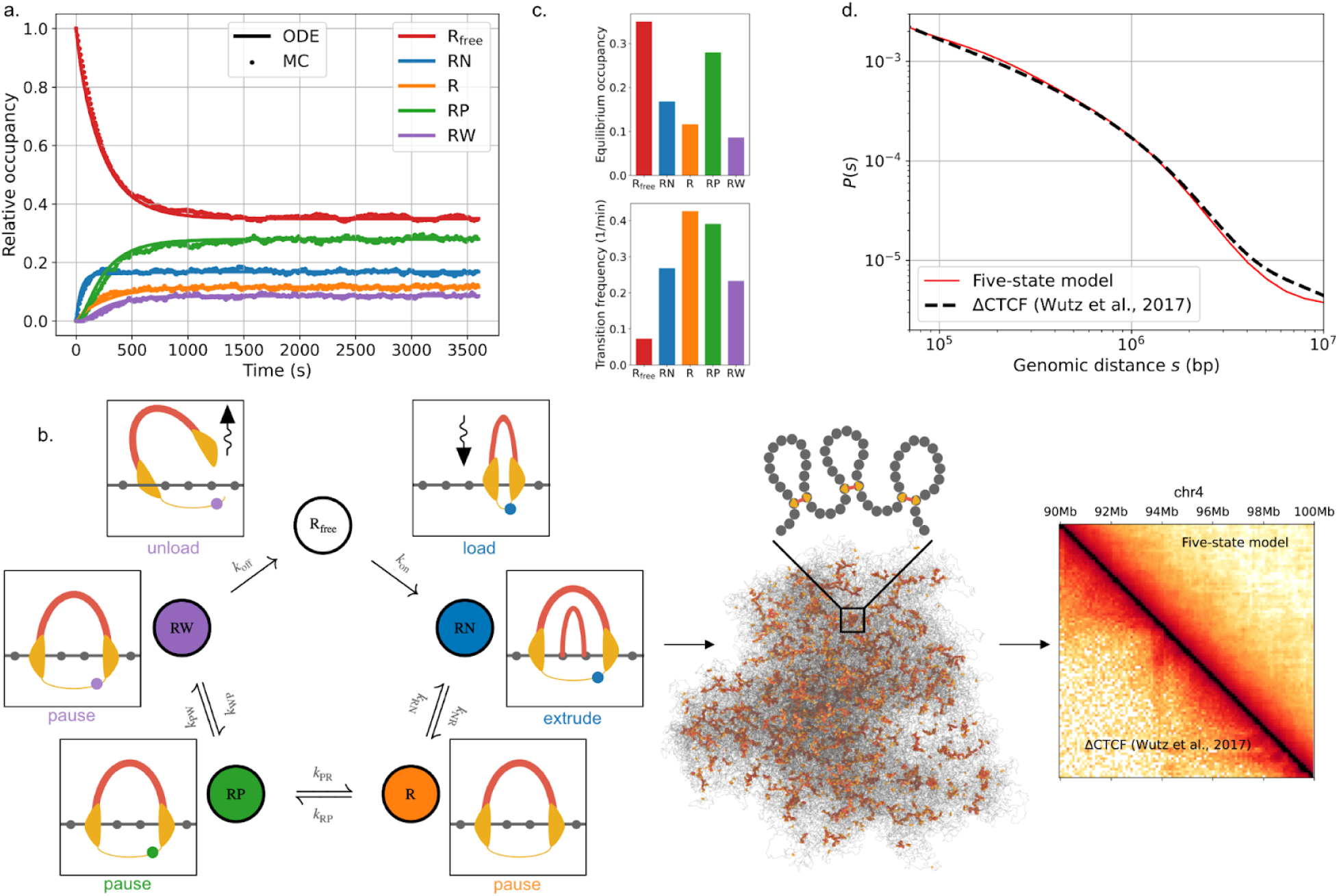
Coupling the cohesin biochemical network with polymer simulations yields quantitative predictions of 3D genome folding. **a.** Model equilibration dynamics, as computed by integration of the mass-action ordinary differential equation (ODE, solid) or by a discrete-time kinetic Monte Carlo (MC, dashed) approach, starting from a fully-unloaded cohesin population (*R_fred_* = 1; see **Methods**). **b.** Schematic representation of the coupled simulation workflow. Extrusion dynamics are computed based on a lattice-based MC scheme (***left***) and used as input for 3D molecular dynamics (***center***) simulations, from which *in silico* contact maps (***right***) or microscopy images may be generated (**Methods**). Transition rates between states in the lattice simulations are parametrized using the five-state reaction network (**Table S1**). Extrusion updates occur only in the NIPBL-bound (RN) state. Unloading happens upon transitioning between the WAPL-bound (RW) and free states, and loading happens upon transitioning between the free and NIPBL-bound states. **c. *Top:*** Equilibrium occupancies, normalized to the total cohesin nuclear content. ***Bottom:*** State transition frequencies, defined as the average number of transitions into each state per cohesin binding window divided by the cohesin residence time (as in (Barth et al., 2023)). **d.** Contact frequency versus distance scaling curves, *P(s)*, either predicted by the five-state cohesin model (red) or obtained from experimental Hi-C in CTCF-depleted HeLa cells (dashed black, (Wutz et al., 2017)).

Using the five-state lattice model, we simulated the dynamics of ∼6,700 five-state extruders on 500 Mb of chromatin at 2.5kb resolution (**Methods**). These values yield a mean density of ∼8.7 loaded cohesins per Mb, consistent with the numbers reported in G1 HeLa cells. To parameterize the average extrusion rate in simulations, which depends on the fraction of time a chromatin-associated extruder is in the NIPBL-bound state (**Fig. 2c**), we used the rate estimated *in vitro* (*v* = 1 *kb*/*s* (Davidson et al., 2019; Kim et al., 2019)). We then obtained *in silico* Hi-C data by generating an ensemble of 5,000 chromatin conformations and recording chromatin contacts (**Fig. 2b**). The resulting contact-versus-distance (*P*(*s*)) scaling curve obtained from simulations quantitatively matched measurements of CTCF-depleted HeLa cells (*R*^2^ > 0. 99, **Fig. 2d**), which provided an optimal reference system for comparison as the current model did not include extrusion barriers. If we instead treated extrusion rate as an adjustable parameter, we found optimal agreement at rates *v* = 850 *bp*/*s* (**Fig. S3**), thus evidencing the physiological relevance of the range reported *in vitro* for human cohesin on both naked DNA and chromatinized templates (*v* ∼ 0. 5 − 1*kb*/*s* (Davidson et al., 2019; Kim et al., 2019)).

The *P(s)* curve predicted by the five-state model was quantitatively similar to those from one- and two-state models of extrusion — which respectively assume a constant extrusion rate with or without immediate rebinding upon unloading ((Buckle et al., 2018; Coßmann et al., 2023; Goloborodko et al., 2016)**, Fig. S4a**). Likely because Hi-C is a population-level analysis, this metric does not reflect the additional heterogeneity in single-extruder properties provided by the five-state model (**Fig. S4b-e; Movie M1**). In contrast, extruder heterogeneity is required to reproduce the dispersion observed *in vitro* for single-molecule properties like the extrusion rate (Davidson et al., 2019; Kim et al., 2019), alongside their association with cohesin biochemical state – as recently reported for NIPBL binding events (Barth et al., 2023). Indeed, the state occupancies and transitions predicted by the five-state model are consistent with multiple experimental observations. First, only a small fraction of all interphase cohesin complexes are predicted to be actively extruding (17%, **Fig. 2c**), quite close to *in vitro* measurements in standard buffer conditions (∼18%, (Pobegalov et al., 2023)). Second, loaded cohesins transition into the actively-extruding state about 0.3 times per minute on average, close to the rate suggested by recent *in vitro* experiments (∼1 per minute (Barth et al., 2023), **Methods**).

Collectively, our coupled five-state reaction network and polymer model quantitatively predicts extrusion kinetics and resulting 3D genome structure based solely on biophysical measurements, without relying on any input from Hi-C data.

### Five-state cohesin model maps regulator abundance to extrusion dynamics

To quantify how protein levels influence cohesin loop extrusion properties, we performed *in silico* depletions of individual regulatory proteins. We first considered predictions for cohesin residence time after 90% depletion of different cohesin regulators, corresponding to the experimental levels of WAPL or PDS5 after RNAi (Wutz et al., 2017). At this depletion level, the five-state model predicted that cohesin residence time would increase eight-fold after ΔWAPL or two-fold after ΔPDS5 (**Fig. 3a**). These predictions agreed with experimental FRAP after either WAPL or PDS5A+B depletion by RNAi (Wutz et al., 2017). Importantly, cohesin residence times after regulator depletions were not used to fit the five-state model, and thus provided orthogonal model validation. We next considered predictions for the chromatin-bound fraction of cohesin, again after 90% depletion. The five-state model predicted a greater increase after ΔWAPL than ΔPDS5 (∼40% vs. ∼20%, **Fig. 3b**). These values are ordered identically, though slightly lower than experimental estimates (∼65% vs. ∼55%, respectively (Wutz et al., 2017)). The discrepancy for the bound fractions can be alleviated by considering an alternative model, where WAPL and PDS5 simultaneously co-bind RAD21 (Ouyang et al., 2013), and are jointly required for cohesin unloading. However, predicted increases in residence times for this strict co-binding model substantially overestimated those observed in experimental data (**Fig. S5**). In contrast to the increased bound fraction after ΔWAPL or ΔPDS5, the five-state model predicted that NIPBL depletion would decrease the loaded cohesin fraction, consistent with the 45% reduction reported experimentally (Wutz et al. 2017). Since RNAi efficiency was not reported for NIPBL, we considered a range of depletion levels *in silico* and found the best agreement with experiments occurred at 60% depletion.

**Figure 3:**
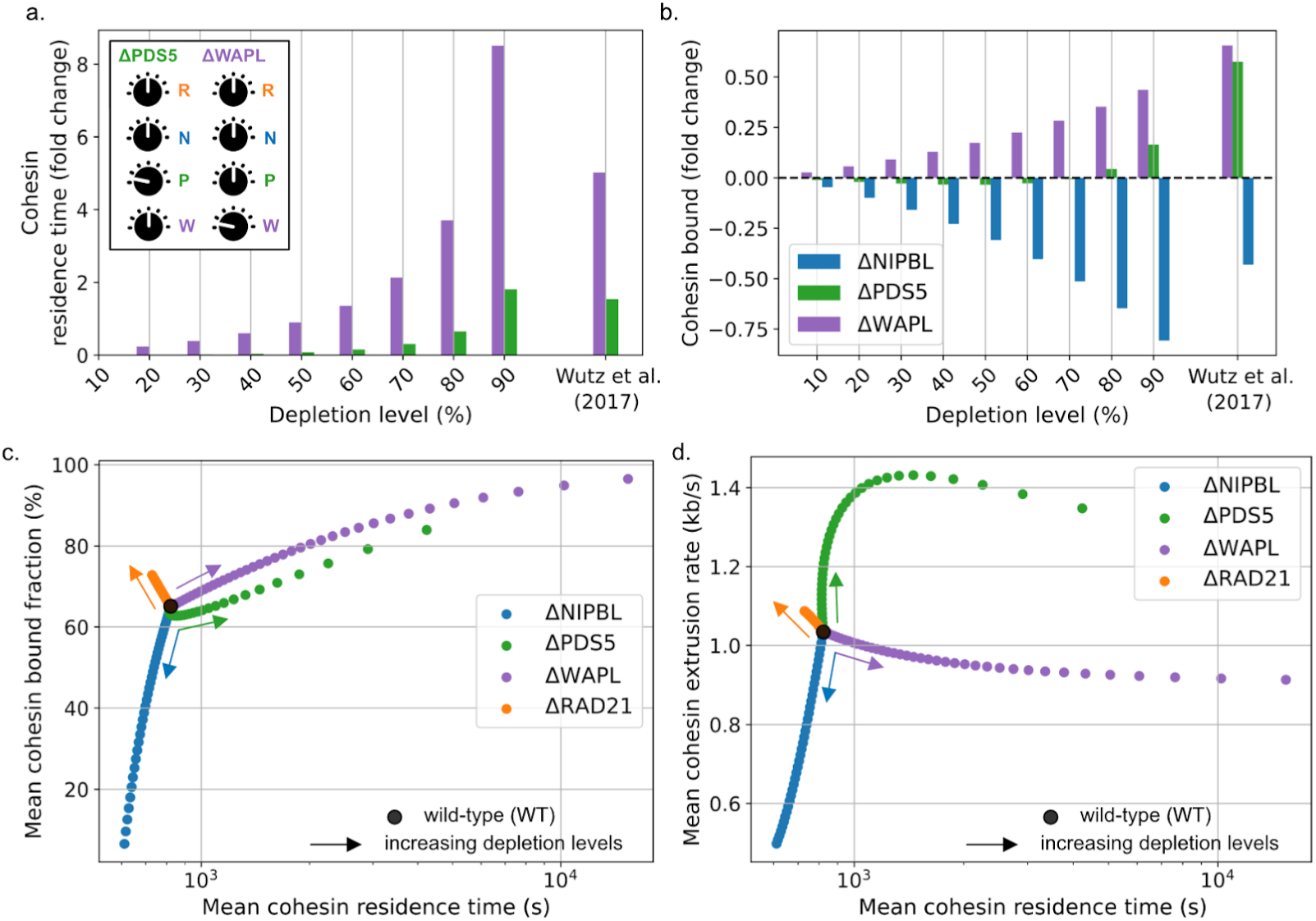
Five-state model predicts non-linear relationship between regulator abundance and extrusion parameters. **a.** Change in cohesin residence time as a function of depletion level for WAPL (purple) and PDS5 (green) in simulations (from 10% to 90%) and experimental data (Wutz et al., 2017). **b.** Change in cohesin bound fraction for simulations and experimental data. **c.** Residence time versus fraction bound as a function of depletion level from the WT values (black circle). Each dot represents a distinct simulated depletion level of the indicated protein in 2.5% increments; arrows show direction of increasing depletion. **d.** Extrusion rate versus residence time, with dots and arrows as in (**c**). Average rate is calculated from the time spent in the NIPBL co-bound state relative to the full cohesin residence time (**Methods**).

More broadly, the five-state model predicted that depletion of individual cohesin regulators could produce non-linear impacts on multiple extrusion properties. We found: (i) ΔNIPBL increased extrusion rate and lowered the bound fraction; (ii) ΔWAPL increased bound fraction and lifetime, (iii) ΔPDS5 changed all three extrusion properties, and (iv) ΔRAD21 chiefly lowered the number of cohesins on chromatin (**Figs. 3c-d)**. Consistent with the ΔNIPBL prediction for a limited decrease in residence time, a minimal reduction in loop lifetimes was reported *in vitro* upon lowering the ratio of NIPBL to RAD21 (Barth et al., 2023). The lower extrusion rate predicted in ΔNIPBL mirrors experimental observations of lower ATP hydrolysis rates by cohesin when the availability of NIPBL is reduced (Davidson et al., 2019; Murayama and Uhlmann, 2014; Petela et al., 2018). Conversely, the higher extrusion rate predicted for ΔPDS5 resulted from a higher frequency of re-binding NIPBL from the bare loaded cohesin state, leading to an upturn in the NIPBL-bound cohesin population (RN), which similarly aligns with experimental observations (Ouyang et al., 2013; van Ruiten et al., 2022). In our model, an increased extrusion rate is eventually compensated by an increased bound cohesin fraction at higher depletion levels of PDS5 (**Fig. 3c**), which also lowers the availability of NIPBL per loaded cohesin. These competing effects lead to a non-monotonic dependence of extrusion rate on PDS5 levels, characterized by an initial increase followed by a moderate decrease at very high PDS5 depletion levels (**Fig. 3c**).

The five-state model also predicts the residence time and bound fraction of NIPBL, PDS5 and WAPL after depletion of any cohesin regulator. For instance, it suggests that ΔWAPL increases the chromatin-associated fraction of NIPBL, yet minimally alters its residence time (**Table S2**) — in agreement with FRAP experiments after perturbation *in vivo* (Rhodes et al., 2017). Similarly, it predicts the chromatin residence time of NIPBL is largely independent of its expression level, consistent with single-molecule observations of cohesin extrusion *in vitro* (Barth et al., 2023). These results highlight how our model captures the coupled dynamics of cohesin regulators beyond their effects on cohesin extrusion.

To explore the synergistic or antagonistic roles of cohesin regulators, we further considered some of their combined depletions *in silico*. The five-state model predicted that the stoichiometric co-depletion of WAPL and NIPBL lead to cohesin loop patterns quantitatively similar to those observed in unperturbed cells, albeit with much longer residence times and slower extrusion rates for individual extruders (**Fig. S6a-d**). Conversely, the co-depletion of WAPL and RAD21 failed to recover the wild-type phenotype, as the increased cohesin residence time induced by ΔWAPL is in this case not offset by a reduction in the rate of extrusion – resulting in a considerable increase in loop sizes (**Fig. S6e-f**). This predicted WAPL/NIPBL compensation yields a mechanistic explanation for the reported rescue of genome folding following combined depletion of WAPL and NIPBL in Hap1 (Haarhuis et al., 2017) and HCT116 cells (Luppino et al., 2022) and illustrates how the interplay between regulators dictates the fine balance between cohesin loading, unloading, and extrusion activity.

Together, the agreement between *in silico* depletions and experimental observations evidence the power of the five-state model to predict how cohesin extrusion dynamics are modulated by variations in regulator abundance.

### Five-state cohesin model predicts consequences of protein depletions on genome conformation

To investigate how altered loop extrusion properties translate into changes in 3D genome folding, we repeated our *in silico* mutant analysis using polymer simulations coupled to the five-state model and computed observables that could be compared with either experimental Hi-C or microscopy. We first investigated the effects of regulator depletions on 3D chromosome morphology by performing *in silico* microscopy for both cohesin and chromatin. We extracted the spatial positions of cohesins and chromatin from individual conformations, rasterized into a 3D voxel grid, and performed convolution with a gaussian kernel (**Fig. 4a**). Visually, as WAPL depletion increases, a granular cohesin backbone emerges, which results from an accumulation of collided extruders along the chromatin fiber (**Fig. 4b**). We quantified this using a “vermicelli score” for the degree of co-localization of chromatin around potential cohesin clusters (**Figs. 4a-c**). This vermicelli score increases with WAPL or PDS5 depletion, but decreases in ΔRAD21 or ΔNIPBL – whose depletion did not prompt vermicelli patterns (**Figs. 4c; S7**). Model predictions agree with experimental *in situ* fluorescence microscopy, which reported vermicelli upon WAPL or PDS5A+B depletion in HeLa, contrasted by a loss of the chromatin-associated cohesin in ΔNIPBL (Wutz et al., 2017).

**Figure 4:**
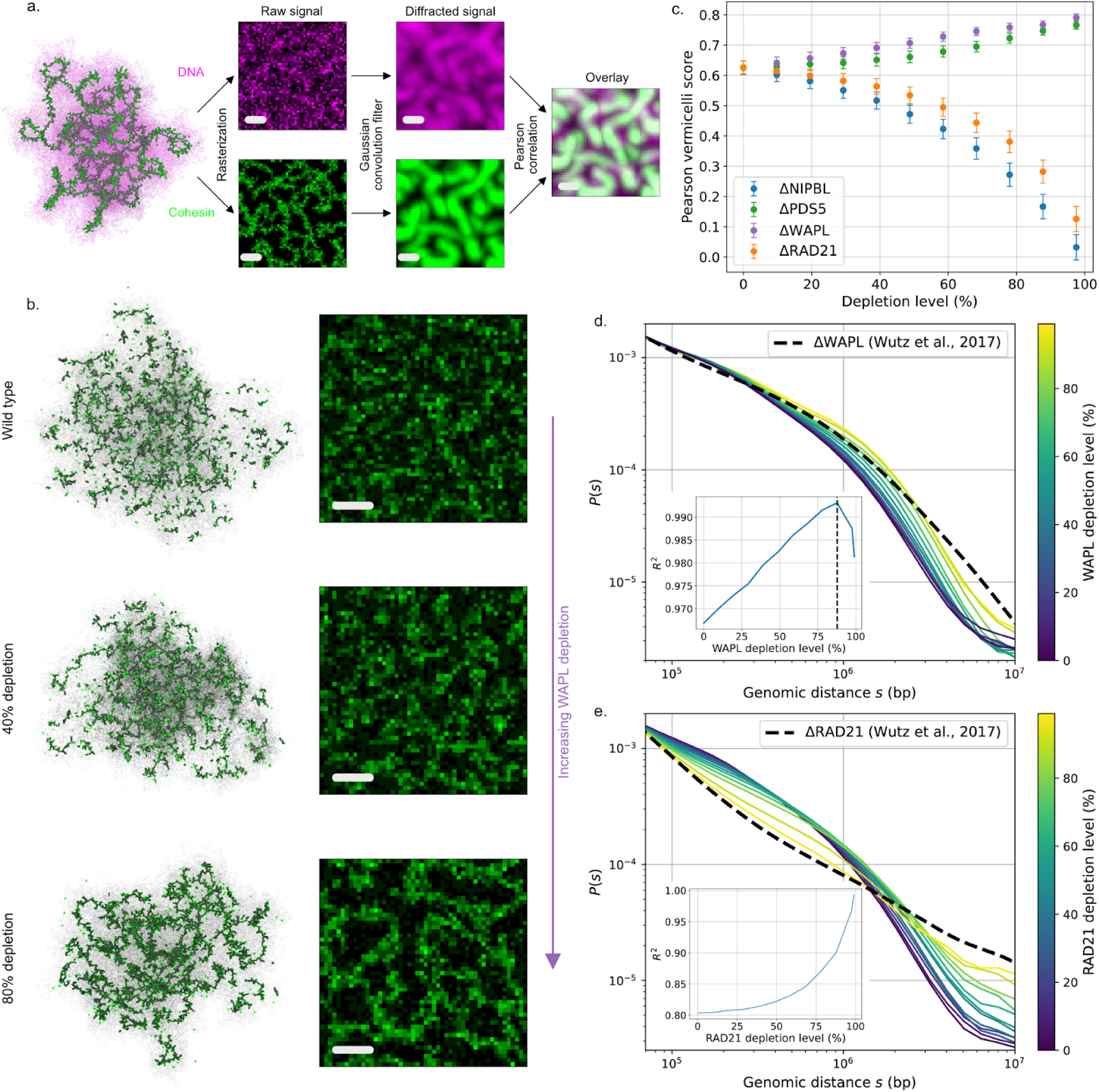
Five-state model maps protein abundance to 3D genome morphology. **a.** Vermicelli score computation workflow. Cohesin (green) and DNA (magenta) spatial positions are separately tagged and binned into discrete 3D voxels. The two resulting rasters are subsequently run through a Gaussian convolution filter to mimic the effects of optical diffraction. The vermicelli score is then defined as the Pearson correlation of the processed cohesin and DNA signal (**Methods**). Scale: 1 µ*m*. **b. *Left:*** Polymer conformations, showing extruder positions (green). ***Right:*** simulated microscopy of RAD21 localization. Both are displayed as a function of WAPL depletion level, showing the emergence of vermicelli. **c.** Vermicelli scores as a function of simulated depletion level for indicated factors. Depletion of WAPL and PDS5 promote vermicelli formation; NIPBL and RAD21 depletion do not. **d.** Contact frequency versus distance curves for simulated ΔWAPL (colored by depletion level), with experimental RNAi depletion (dashed line) from (Wutz et al., 2017). Inset shows best fit is reached at 88% depletion. **e.** Same as **d** for simulated ΔRAD21 and experimental auxin-induced degron (dashed line) from (Wutz et al., 2017). Inset shows best fit is approached at the highest depletion levels considered in simulations (99%).

Auxiliary measures of vermicelli formation, namely the coverage by loops or the collided fraction of loop extruders, largely mirrored the microscopy-based Pearson vermicelli score (**Figs. S8a-b)**. In addition to these intuitive metrics, we also quantified vermicelli formation via a percolation score, computed as the fraction of loaded cohesins comprising the largest cluster of collided extruders. Visually, this score appears to better capture the extent of vermicelli formation, and displays a sharp increase upon depletion of WAPL and PDS5 beyond 80% (**Figs. S7; S8c)**. We note, however, that these alternative quantities are not currently accessible to direct experimental measurements.

To further assay the effects of altered loop extrusion on 3D genome conformations, we extracted *P*(*s*) from *in silico* and *in vivo* Hi-C (**Figs. 4d-e; S9**). We observed that WAPL depletion leads to a rightwards shift of the characteristic “shoulder” in these curves associated with loop extrusion (Fudenberg et al., 2017; Polovnikov et al., 2023). By computing the goodness-of-fit (*R*^2^, **Methods**), the experimental data was best reproduced by WAPL depletion levels of around ∼85% depletion (**Fig. 4d)**, consistent with the RNAi efficiency estimated experimentally (Wutz et al., 2017). PDS5 depletion displayed a similar shift in the shoulder, again congruent with experimental data, albeit without evidence for a clearly best-fitting degradation level (**Fig. S9a**). Conversely, ΔRAD21 had an entirely different impact on the *P*(*s*) in both simulation and experiments, and instead resulted in the gradual disappearance of the shoulder (**Fig. 4e**). In this case, optimal agreement occurred at RAD21 depletion levels greater than 99%, consistent with its highly efficient removal via auxin-induced degradation (Wutz et al., 2017). Simulated ΔNIPBL also led to a gradual disappearance of the shoulder, but the corresponding experimental Hi-C data in NIPBL-depleted HeLa cells have, to our knowledge, yet to be reported.

Collectively, our *in silico* microscopy and Hi-C results indicate that the minimalistic description of cohesin biochemistry provided by the five-state model nevertheless captures how the levels of cohesin and its regulators jointly determine genome folding.

## Discussion

In summary, we present a quantitative model of the cohesin biochemistry underlying interphase loop extrusion, parametrized solely by biophysical data on the dynamics of cohesin and its regulators. We find that a unique minimal reaction network featuring five cohesin states can be fully reconciled with protein abundances, FRAP, and cohesin IP assays after individual regulator depletions in HeLa cells. Combining this five-state model with polymer simulations enables direct predictions of 3D genome folding from nuclear protein levels, and evidences how the transient binding of cohesin regulators may modulate loop extrusion kinetics across multiple time- and length-scales.

The five-state model concisely articulates knowledge of cohesin biochemistry and the respective functions of cohesin regulators. The loading action of NIPBL (Ciosk et al., 2000) and the unloading function of WAPL (Gandhi et al., 2006) have both been long-hypothesized based on sister chromatid cohesion phenotypes. The contributions of PDS5 appear more multifaceted, and have been proposed to include both the promotion of cohesin unloading in cooperation with WAPL (Shintomi and Hirano, 2009) as well as competition with NIPBL for the same RAD21 binding site (Kikuchi et al., 2016). These functions naturally emerge from the five-state model, whose network topology suggests that PDS5 plays a role not only in the recruitment of WAPL, but also hinders the re-binding of NIPBL to loaded cohesin. Additionally, our modeling approach enabled us to rule out large swaths of incompatible alternate topologies (**Fig. S1, S2**) and provide mathematical support for the concept that distinct loading and unloading pathways operate for loop extrusion as well as sister chromatid cohesion.

The five-state model also provides mechanistic insight into biophysical and genomic observations. First, the five-state model demonstrates how the relatively rapid exchange of cohesin regulators can regulate loop extrusion dynamics over much longer time-scales. For instance, it quantitatively reconciles the short residence time of WAPL (∼1 min, (Ladurner et al., 2016)) with the dramatic increase in cohesin residence time induced by its depletion (>1 hour, (Tedeschi et al., 2013)), as well as the associated emergence of the vermicelli morphology at the chromosome-wide level (Tedeschi et al., 2013; Wutz et al., 2017). Furthermore, our model argues that contrasting conclusions drawn either from RNAi or more efficient auxin-induced degradation (de Wit and Nora, 2023) could result simply from the different degrees of depletion achieved by the different experimental protocols, combined with the predicted nonlinear relationships between protein levels and extrusion properties.

Beyond the kinetics of the core complex, the five-state model highlights informative new experiments based on monitoring the dynamics of regulators in various conditions. Separate measurements for PDS5A and PDS5B binding kinetics could enable us to refine model predictions, and may help resolve the moderate discrepancies in cohesin properties and contact-versus-distance scaling in simulated ΔPDS5. Existing experiments do not exclude a model where PDS5 and WAPL simultaneously bind RAD21 and are jointly required for cohesin unloading (Ouyang et al., 2013; Ouyang and Yu, 2017). Our modeling indicates, however, that measurements of the PDS5 residence time before and after WAPL depletion could support or exclude this possibility: if the PDS5 lifetime does not substantially change in ΔWAPL cells, this would support a model where cohesin is unloaded via a co-bound PDS5/WAPL state (**Fig. S5: Table S3**). Conversely, an increase of the PDS5 residence time upon WAPL depletion would suggest that the role of PDS5 primarily lies in the recruitment of WAPL, which in turn acts as the standalone primary cohesin unloader (**Table S2**). Such measurements of regulator kinetics in perturbed cells are currently largely lacking, but would now provide important benchmarks due to the powerful predictive ability of biochemical reaction network models.

By virtue of its minimalist approach, our five-state model provides an ideal basis for the inclusion of additional details of cohesin biochemistry in future work. More complex models could incorporate other known regulators of cohesin such as SA1/2, SCC4 and Sororin (Pezic et al., 2017), as well as the roles of CTCF (van Ruiten and Rowland, 2021) and SMC3 acetylation (Cuadrado and Losada, 2020; Wutz et al., 2020). Accounting for these regulators will be crucial for future models of cohesin biochemistry throughout the cell cycle, including for the establishment of sister chromatid cohesion (Peters and Nishiyama, 2012). The implementation of more complex network topologies with additional transitions will also be of future interest. This includes models with transient, NIPBL-independent association with chromatin (Haarhuis et al., 2017; Ladurner et al., 2016) and WAPL-independent cohesin dissociation (Srinivasan et al., 2019). Consideration of such pathways could help refine model predictions — e.g., by providing a background population of loaded cohesins upon full NIPBL removal for the former, or by imposing a finite cohesin residence time on chromatin upon full WAPL knockout for the latter.

Since the details of how nanoscale conformational changes result in loop extrusion remain uncertain (Dekker et al., 2023), we assumed that individual extruders symmetrically reel in chromatin. Many alternatives and elaborations are likely, including: asymmetric extrusion with switching (Banigan et al., 2020; Barth et al., 2023), a dependence of the loop extrusion rate on chromatin tension (Nomidis et al., 2022) and/or local conformation (Conforto et al., 2024), cohesin backtracking and bypassing (Banigan and Mirny, 2020) or capture of spatially proximal chromatin *in trans* (Bonato and Michieletto, 2021). Future models will be required to consider locus-specific cohesin properties such as targeted loading or the interplay between cohesin and the transcription machinery (Corsi et al., 2023), which could differentially modulate the extrusion rate in passive and actively-transcribed regions (Banigan et al., 2023). Nonetheless, five-state extruders with simple “blocking” collisions that operate uniformly across the genome produce contact frequency versus distance curves in excellent agreement with experimental Hi-C. In HeLa cells, our results thus argue that cohesin kinetics and collisions are the dominant factors determining interphase loop sizes, to which additional mechanisms potentially contribute as higher-order effects. Similar conclusions were recently reported for condensin-based loop extrusion in mitotic cells (Samejima et al., 2024), which suggests that collision-based encounter rules between extruders of the same type could be evolutionarily-conserved across SMC complexes.

To conclude, the five-state model establishes a minimalistic description of cohesin biochemistry capable of quantitatively capturing the roles of key cohesin regulators. It puts forth a molecular paradigm for interphase loop extrusion centered on cohesin as a multi-state motor. Our results highlight the ability of simple biophysical models to integrate data from multiple orthogonal modalities, including *in vitro* motor assays, quantitative *in vivo* measurements, genomics, and *in situ* immunofluorescence microscopy to build a holistic picture of chromosome organization. Altogether, our framework illuminates how cells can harness loop extrusion by fine tuning regulator abundances across cell types and states, and bridges the gap between our molecular and genome-scale understanding of chromosome organization.

## Methods

### Literature curation of the abundance and dynamics of cohesin and its regulators

Determining the transition rates for our minimal model of interphase cohesin chemistry requires three quantities – namely, abundance, bound fraction, and residence times measured in unperturbed cells – for each regulator considered. For a literature estimate of abundance, we averaged the cohesin regulator numbers quantified in HeLa cells by mass spectrometry (Bekker-Jensen et al., 2017) and fluorescence correlation spectroscopy (FCS) (Holzmann et al. 2019). Absolute protein copy numbers were converted to genomic densities by considering a ∼ 19. 539 *Gb* average genome size, as previously reported for diploid HeLa cells (Kesel et al., 2016). Since FCS measurements of PDS5 abundance are to our knowledge currently lacking, we used as an alternative estimate the mean stoichiometric PDSA/B-to-WAPL ratio reported in HeLa immunoprecipitation assays via SMC1 and SMC3 pulldowns (Ding et al., 2011). We similarly curated Fluorescence Recovery After Photobleaching (FRAP) data to obtain bound fractions and residence times for cohesin regulators in HeLa cells from the following publications: RAD21 from Hozmann et al. (Holzmann et al. 2019); NIPBL from Rhodes et al., (Rhodes et al., 2017); WAPL and PDS5 from Ladurner et al., (Ladurner et al., 2016). These values (**Table 1**) were employed to ascertain the reaction network of unperturbed HeLa cells.

### Assumptions for building a minimal cohesin biochemical reaction network

1. RAD21 may be taken as a proxy for the core cohesin complex:

○ Based on structural insights that RAD21 acts as a “nexus” for the recruitment of cohesin regulatory factors NIPBL, PDS5, and WAPL (Muir et al., 2016).
○ The dynamic residence time of RAD21, SMC3, SMC1, and SA1 are all similarly in the tens of minutes range (Gerlich et al., 2006; Kueng et al., 2006; Wutz et al., 2020).
2. Cohesin regulators do not bind chromatin in the absence of RAD21:

○ NIPBL: (Rhodes et al., 2017) reports that RAD21 depletion releases the majority of (but not all) NIPBL from chromatin, suggesting that RAD21 association is the dominant pathway for loading NIPBL onto chromosomes. This is also consistent with the disappearance of NIPBL-associated ChIP-seq peaks upon removal of cohesin (Banigan et al., 2023).
○ WAPL: Kueng et al. 2006 reports that WAPL cannot be detected on chromatin in RAD21-depleted cells.
○ PDS5: Panizza et al. 2000 reports that the recruitment of PDS5 of chromatin is drastically reduced in cells not expressing RAD21. This is also consistent with reports that disrupting the RAD21-PDS5 binding interface largely abolishes the association of PDS5 with chromatin (Muir et al., 2016).
3. Co-bound states involving the simultaneous association of multiple regulators are neglected:

○ PDS5+NIPBL: Reports that PDS5 competes with NIPBL for RAD21 binding (Kikuchi et al., 2016; Petela et al., 2018) support this hypothesis.
○ WAPL+NIPBL: Supported by Arruda, Bryan, and Dowen 2022, which report a small (but non-zero) co-bound population in WAPL-NIPBL co-IP experiments.
○ WAPL+PDS5: Although WAPL, RAD21 and PDS5 have been recently suggested to be able to form a tripartite complex using AlphaFold (Nasmyth et al., 2023), experimental observations in reconstituted protein assays have revealed that WAPL may also stably associate with RAD21 in the absence of PDS5 (Gandhi et al., 2006). Similarly, the chromatin association of PDS5 *in vivo* was largely unaffected by WAPL depletion (Chan et al., 2013) arguing against a substantial population of WAPL and PDS5 being concomitantly bound to cohesin, despite a reported role for PDS5 in the recruitment of WAPL (Chan et al., 2012)*. In vitro* studies have further reported conflicting evidence of direct interactions between WAPL and PDS5 (Muir et al., 2016; Ouyang et al., 2013), although co-IP experiments support the existence of WAPL-PDS5A co-binding in presence of RAD21 (Gandhi et al., 2006; Kueng et al., 2006). In light of this uncertainty, we neglect the simultaneous binding of PDS5 and WAPL onto RAD21 as a first approximation. However, we also explore a model with a strictly co-bound WAPL+PDS5 state and describe an experimental signature to unambiguously assess its relevance.
4. The A and B paralogs of PDS5 may be amalgamated:

○ First, the two paralogs associate with cohesin in a mutually-exclusive fashion (Losada et al., 2005). Second, studies have suggested that PDS5A/B share a large structural and functional redundancy (Zhang et al., 2009). Further investigations would be required to clarify their individual roles and potential differential effects on extrusion (Arruda et al., 2022; Zhang et al., 2021) distinct FRAP measurements for PDS5A and PDS5B are, to our knowledge, currently lacking.
5. We consider models with 8 non-zero transition rates and four loaded states:

○ This precludes the study of potentially interesting models with additional transitions, e.g. transient binding of non-extrusive RAD21 is neglected (Ladurner et al., 2016), along with potential WAPL-independent cohesin dissociation (Srinivasan et al., 2019).

### Rate mapping

Cohesin state transition rates were inferred from the experimentally-measured bound fractions and residence times of individual proteins. Assuming mass action kinetics, we derived a system of coupled ordinary differential equations (ODEs) for the populations of RAD21, NIPBL, PDS5 and WAPL. This system of ODEs involves 8 unknown cohesin state transition rates (**Fig. 1**). Accordingly, we derived 8 mathematical constraints from the chromatin residence time and bound fraction for each molecular species. Combining these, we arrive at a system of 8 linear equations, which may be solved symbolically to obtain explicit expressions for the transition rates as rational functions of the protein absolute abundances, residence times, and bound fractions (**Table 1**; **Supp. Sec. I**).

### Lattice model for multi-state extrusion kinetics

We simulated loop extrusion as a discrete-time process on a 1D lattice with timestep τ_1*D*_ at a genomic resolution of *l* ∼ 2. 5 *kb* per site. When extruders are loaded, a left and right leg are placed on adjacent lattice sites. During each update step, a given extruder may be stochastically loaded, unloaded, or transition into a new loaded state within the cohesin biochemical network (*RN*, *R*, *RP*, *RW*) based on a simple discrete-time kinetic Monte-Carlo sampling of the five-state reaction network. After the possible state update, if an extruder is in the active NIPBL-bound (*RN*) state, each leg of the extruder moves one lattice site outwards, provided that adjacent lattice sites are unoccupied. After each update, the positions of all extruder legs and extruder states are recorded. For simplicity, we assumed that the corresponding transition rates between different cohesin states are uniform across all sites — i.e., that cohesin physico-chemical properties are independent of the local genomic context — and may thus be identically set to the respective values inferred from the rate mapping procedure (**Table S1**).

Denoting by [*RN*] the total numbers of RAD21 molecules bound by NIPBL at equilibrium, the mean extrusion rate at steady state reads as *v* = 2*l*/τ × [*RN*]/[*R*]_*loaded*_, where _1*D*_ [*R*]_*loaded*_ = [*RN*] + [*R*] + [*RP*] + [*RN*] indicates the total equilibrium population of chromatin-associated RAD21 and the factor 2 accounts for the two legs of the cohesin complex. The five-state model predicts the value of the active-to-loaded extruder ratio as [*RN*]/[*R*]_*loaded*_ ≃ 26 % in wild-type HeLa cells. Using the typical extrusion rate of *v* ∼ 1 *kb*/*s* estimated for cohesin by single-molecule imaging *in vitro* (Davidson et al., 2019; Kim et al., 2019), we may thus infer that each lattice step τ_1*D*_ corresponds to approximately ∼ 1. 25 *s* of physical time.

Using these values, the model predicts a transition frequency into the NIPBL-bound state of about 0.3 times per minute (**Fig. 2c**). This rate is slightly slower than an experimental value of ∼1/min, as inferred from the ∼0.5/min rate of cohesin direction changes reported in single-molecule *in vitro* assays (Barth et al., 2023), where the factor of 2 accounts for the two possible extrusion directions after each NIPBL binding event. Note that since extrusion traces lacking a direction switch were excluded from analysis in (Barth et al., 2023), their estimated experimental rate of ∼1/min likely provides an upper bound for the average transition frequency into the NIPBL-associated state.

### Polymer model of multi-state extrusion

We model a 500 Mb-long chromatin region as a linear polymer comprising 200,000 monomers of 2. 5 *kb* each, such that each site from the lattice model corresponds to a unique individual bead in the chromatin chain. As in previous investigations (Nuebler et al., 2018), we use a spatial extent of each bead of *b* ∼ 50 *nm*, consistent with recent estimates of chromatin compaction of 50 *kb*/µ*m* in eukaryotes (budding yeast, (Arbona et al., 2017)). To model the impact of extruders on 3D polymer conformations, we generated extruder positions using the 1D lattice model and created additional bonds between pairs of monomers occupied by the two legs of each extruder. Polymer simulations were run using the the polychrom-hoomd package (https://github.com/open2c/polychrom-hoomd), based on the HooMD molecular dynamics engine (Anderson et al., 2020), considering a 20% polymer volume fraction combined with a polynomial soft excluded-volume potential (Gabriele et al., 2022) (see https://github.com/fudenberg-research-group/five_state_cohesin and https://github.com/open2c/polychrom-hoomd for full implementation details). This concentration amounts to an approximate chromatin density of ^3^, consistent with the typical value obtained using the average HeLa genome size (19. 539 *Gb*, (Kesel et al., 2016)) and nuclear diameter (15. 7 µ*m*, (James and Giorgio, 2000)), assuming a spherical nuclear shape. Numerical integration was performed using dissipative particle dynamics (DPD) with a dimensionless 3D integration timestep τ_3*D*_ = 0. 005 (Phillips et al., 2011). At long times (≽ 100 *s*), the computed (Rouse) diffusion coefficient of individual monomers reads as 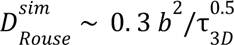 (**Fig. S3a**). The mapping of simulation to experimental times is performed by matching 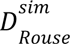 to the experimental value 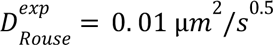 estimated in yeast chromatin (Hajjoul et al., 2013), which yields τ_3*D*_ ∼ 5 *ms*. Alternatively, the presence of cohesin with wild-type extrusion parameters (**Table 1**) leads to a simulated super-Rousean anomalous diffusion coefficient 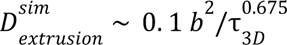 at short times (≼ 100 *s*), which may be compared to the experimental value 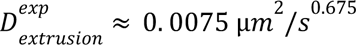 estimated in CTCF-depleted mESCs over a similar time interval (Mach et al., 2022) **(Fig. S3a)**. This procedure similarly leads to τ_3*D*_ ∼ 5 *ms*, which evidences the relative robustness of this time mapping. The number of MD steps performed between each extruder update is then given by *N*_3*D*/1*D*_ = τ_1*D*_/τ_3*D*_ = 250. To assess the sensitivity of our predictions to this number, we analyzed the mean-squared error in the simulated contact frequency versus distance curve (*P*(*s*); see below), compared to the experimental Hi-C profile observed in CTCF-depleted HeLa cells (Wutz et al. 2017), across 10 different *N*_3*D*/1*D*_ values in the range [100, 1000] (**Fig. S3b-c**). This procedure yields an optimal agreement between theory and experiment for *N*_3*D*/1*D*_ = 300, corresponding to a mean extrusion rate of *v* ∼ 850 *bp*/*s* – highly consistent with biophysically-inferred parameters (*N*_3*D*/1*D*_ = 250, *v* ∼ 1 *kb*/*s*).

### Contact frequency versus distance curves

For the calculation of contact frequency versus distance (*P*(*s*)) curves, we used the monomerResolutionContactMapSubchains function from the contact_maps module as implemented in the polykit package (https://github.com/open2c/polykit). Contact maps were computed at the single monomer level using a default cutoff distance *R*_*c*_ = 2. 3 *b* ∼ 115 *nm*, consistent with the typical capture radius assumed in standard Hi-C experiments (McCord et al., 2020). *P*(*s*) scaling curves were then directly computed from the maps using the expected_cis function of the cooltools library (Abdennur et al., 2024). To obtain the experimental contact frequency distance curves, *P_exp_*(*s*), we re-processed experimental Hi-C datasets from HeLa cells (Wutz et al. 2017) using the distiller pipeline (https://github.com/open2c/distiller-nf, (Goloborodko et al., 2022)), extracting contacts with pairtools (https://github.com/open2c/pairtools, (Open2C et al., 2024)), and binning to 10kb resolution with cooler (https://github.com/open2c/cooler, (Abdennur and Mirny, 2020)). To quantify agreement between simulations and experiment, a goodness-of-fit parameter (*R*^2^) was then defined as

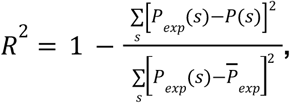

where the summation and average 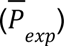 are both performed over the genomic range [50 *kb*, 10 *Mb*] using 4950 log-spaced bins. The lower bound of 50 *kb* was chosen to lie above the threshold of 40 *kb*, beyond which HindIII-based Hi-C data appears unaffected by restriction and ligation artifacts resulting from the experimental library preparation protocol (Naumova et al., 2013).

### Numerical microscopy and Pearson vermicelli score

To matching the typical scanning resolution of confocal microscopes, 3D voxels of dimension 100 *nm* × 100 *nm* × 100 *nm* are used for the binning of cohesin and DNA spatial positions. The resulting cubic rasters feature 50 voxels along each axis, corresponding to a total field of view with linear dimension 5 µ*m*. To mimic the presence of unloaded extruders, additional diffusive cohesins are randomly and uniformly distributed throughout the sample, with numbers matching the unloaded population of RAD21 predicted by the five-state model in each condition. A Gaussian convolution filter with standard deviation of 250 *nm* is subsequently applied to approximate the Airy point spread function. The Pearson correlation score between the diffracted cohesin and DNA signal is finally computed by averaging across 5000 MD frames obtained from 5 independent simulations for each sample.

## Acknowledgements

We thank Elphège Nora, Anton Goloborodko, Erika Anderson, and Gordana Wutz for helpful discussions and detailed feedback. The authors are supported by NIGMS R35GM143116 to GF.

## References

Abdennur, N., Abraham, S., Fudenberg, G., Flyamer, I.M., Galitsyna, A.A., Goloborodko, A., Imakaev, M., Oksuz, B.A., Venev, S.V., Xiao, Y., 2024. Cooltools: Enabling high-resolution Hi-C analysis in Python. PLOS Comput. Biol. 20, e1012067. 10.1371/journal.pcbi.1012067

Abdennur, N., Mirny, L.A., 2020. Cooler: scalable storage for Hi-C data and other genomically labeled arrays. Bioinforma. Oxf. Engl. 36, 311–316. 10.1093/bioinformatics/btz540

Alipour, E., Marko, J.F., 2012. Self-organization of domain structures by DNA-loop-extruding enzymes. Nucleic Acids Res. 40, 11202–11212. 10.1093/nar/gks925

Anderson, J.A., Glaser, J., Glotzer, S.C., 2020. HOOMD-blue: A Python package for high-performance molecular dynamics and hard particle Monte Carlo simulations. Comput. Mater. Sci. 173, 109363. 10.1016/j.commatsci.2019.109363

Arbona, J.-M., Herbert, S., Fabre, E., Zimmer, C., 2017. Inferring the physical properties of yeast chromatin through Bayesian analysis of whole nucleus simulations. Genome Biol. 18, 81. 10.1186/s13059-017-1199-x

Arruda, N.L., Bryan, A.F., Dowen, J.M., 2022. PDS5A and PDS5B differentially affect gene expression without altering cohesin localization across the genome. Epigenetics Chromatin 15, 30. 10.1186/s13072-022-00463-6

Banigan, E.J., Mirny, L.A., 2020. Loop extrusion: theory meets single-molecule experiments. Curr. Opin. Cell Biol. 64, 124–138. 10.1016/j.ceb.2020.04.011

Banigan, E.J., Tang, W., van den Berg, A.A., Stocsits, R.R., Wutz, G., Brandão, H.B., Busslinger, G.A., Peters, J.-M., Mirny, L.A., 2023. Transcription shapes 3D chromatin organization by interacting with loop extrusion. Proc. Natl. Acad. Sci. U. S. A. 120, e2210480120. 10.1073/pnas.2210480120

Banigan, E.J., van den Berg, A.A., Brandão, H.B., Marko, J.F., Mirny, L.A., 2020. Chromosome organization by one-sided and two-sided loop extrusion. eLife 9, e53558. 10.7554/eLife.53558

Barth, R., Davidson, I.F., Torre, J. van der, Taschner, M., Gruber, S., Peters, J.-M., Dekker, C., 2023. SMC motor proteins extrude DNA asymmetrically and contain a direction switch. 10.1101/2023.12.21.572892

Bekker-Jensen, D.B., Kelstrup, C.D., Batth, T.S., Larsen, S.C., Haldrup, C., Bramsen, J.B., Sørensen, K.D., Høyer, S., Ørntoft, T.F., Andersen, C.L., Nielsen, M.L., Olsen, J.V., 2017. An Optimized Shotgun Strategy for the Rapid Generation of Comprehensive Human Proteomes. Cell Syst. 4, 587–599.e4. 10.1016/j.cels.2017.05.009

Bonato, A., Michieletto, D., 2021. Three-dimensional loop extrusion. Biophys. J. 120, 5544–5552. 10.1016/j.bpj.2021.11.015

Brackey, C.A., Marenduzzo, D., Gilbert, N., 2020. Mechanistic modeling of chromatin folding to understand function. Nat. Methods 17, 767–775. 10.1038/s41592-020-0852-6

Buckle, A., Brackley, C.A., Boyle, S., Marenduzzo, D., Gilbert, N., 2018. Polymer Simulations of Heteromorphic Chromatin Predict the 3D Folding of Complex Genomic Loci. Mol. Cell 72, 786–797.e11. 10.1016/j.molcel.2018.09.016

Chan, K.-L., Gligoris, T., Upcher, W., Kato, Y., Shirahige, K., Nasmyth, K., Beckouët, F., 2013. Pds5 promotes and protects cohesin acetylation. Proc. Natl. Acad. Sci. U. S. A. 110, 13020–13025. 10.1073/pnas.1306900110

Chan, K.-L., Roig, M.B., Hu, B., Beckouët, F., Metson, J., Nasmyth, K., 2012. Cohesin’s DNA exit gate is distinct from its entrance gate and is regulated by acetylation. Cell 150, 961–974. 10.1016/j.cell.2012.07.028

Ciosk, R., Shirayama, M., Shevchenko, A., Tanaka, T., Toth, A., Shevchenko, A., Nasmyth, K., 2000. Cohesin’s binding to chromosomes depends on a separate complex consisting of Scc2 and Scc4 proteins. Mol. Cell 5, 243–254. 10.1016/s1097-2765(00)80420-7

Conforto, F., Fosado, Y.A.G., Michieletto, D., 2024. Active Fluidification of Entangled Polymers by Loop Extrusion. 10.48550/arXiv.2401.17232

Corsi, F., Rusch, E., Goloborodko, A., 2023. Loop extrusion rules: the next generation. Curr. Opin. Genet. Dev. 81, 102061. 10.1016/j.gde.2023.102061

Coßmann, J., Kos, P.I., Varamogianni-Mamatsi, V., Assenheimer, D., Bischof, T., Kuhn, T., Vomhof, T., Papantonis, A., Giorgetti, L., Gebhardt, J.C.M., 2023. Increasingly efficient chromatin binding of cohesin and CTCF supports chromatin architecture formation during zebrafish embryogenesis. 10.1101/2023.12.08.570809

Cuadrado, A., Losada, A., 2020. Specialized functions of cohesins STAG1 and STAG2 in 3D genome architecture. Curr. Opin. Genet. Dev. 61, 9–16. 10.1016/j.gde.2020.02.024

Davidson, I.F., Bauer, B., Goetz, D., Tang, W., Wutz, G., Peters, J.-M., 2019. DNA loop extrusion by human cohesin. Science 366, 1338–1345. 10.1126/science.aaz3418

Davidson, I.F., Peters, J.-M., 2021. Genome folding through loop extrusion by SMC complexes. Nat. Rev. Mol. Cell Biol. 22, 445–464. 10.1038/s41580-021-00349-7

de Wit, E., Nora, E.P., 2023. New insights into genome folding by loop extrusion from inducible degron technologies. Nat. Rev. Genet. 24, 73–85. 10.1038/s41576-022-00530-4

Dekker, C., Haering, C.H., Peters, J.-M., Rowland, B.D., 2023. How do molecular motors fold the genome? Science 382, 646–648. 10.1126/science.adi8308

Ding, C., Li, Y., Kim, B.-J., Malovannaya, A., Jung, S.Y., Wang, Y., Qin, J., 2011. Quantitative analysis of cohesin complex stoichiometry and SMC3 modification-dependent protein interactions. J. Proteome Res. 10, 3652–3659. 10.1021/pr2002758

Fudenberg, G., Abdennur, N., Imakaev, M., Goloborodko, A., Mirny, L.A., 2017. Emerging Evidence of Chromosome Folding by Loop Extrusion. Cold Spring Harb. Symp. Quant. Biol. 82, 45–55. 10.1101/sqb.2017.82.034710

Fudenberg, G., Imakaev, M., Lu, C., Goloborodko, A., Abdennur, N., Mirny, L.A., 2016. Formation of Chromosomal Domains by Loop Extrusion. Cell Rep. 15, 2038–2049. 10.1016/j.celrep.2016.04.085

Gabriele, M., Brandão, H.B., Grosse-Holz, S., Jha, A., Dailey, G.M., Cattoglio, C., Hsieh, T.-H.S., Mirny, L., Zechner, C., Hansen, A.S., 2022. Dynamics of CTCF- and cohesin-mediated chromatin looping revealed by live-cell imaging. Science 376, 496–501. 10.1126/science.abn6583

Gandhi, R., Gillespie, P.J., Hirano, T., 2006. Human Wapl is a cohesin-binding protein that promotes sister-chromatid resolution in mitotic prophase. Curr. Biol. CB 16, 2406–2417. 10.1016/j.cub.2006.10.061

Gerlich, D., Koch, B., Dupeux, F., Peters, J.-M., Ellenberg, J., 2006. Live-cell imaging reveals a stable cohesin-chromatin interaction after but not before DNA replication. Curr. Biol. CB 16, 1571–1578. 10.1016/j.cub.2006.06.068

Gibcus, J.H., Samejima, K., Goloborodko, A., Samejima, I., Naumova, N., Nuebler, J., Kanemaki, M.T., Xie, L., Paulson, J.R., Earnshaw, W.C., Mirny, L.A., Dekker, J., 2018. A pathway for mitotic chromosome formation. Science 359, eaao6135. 10.1126/science.aao6135

Golfier, S., Quail, T., Kimura, H., Brugués, J., 2020. Cohesin and condensin extrude DNA loops in a cell cycle-dependent manner. eLife 9, e53885. 10.7554/eLife.53885

Goloborodko, A., Imakaev, M.V., Marko, J.F., Mirny, L., 2016. Compaction and segregation of sister chromatids via active loop extrusion. eLife 5, e14864. 10.7554/eLife.14864

Goloborodko, A., Venev, S.V., Spracklin, G., Abdennur, N., Galitsyna, A.A., Shaytan, A., Flyamer, I.M., Di Tommaso, P., Kolchenko, S., 2022. distiller. 10.5281/zenodo.1490628.

Haarhuis, J.H.I., van der Weide, R.H., Blomen, V.A., Yáñez-Cuna, J.O., Amendola, M., van Ruiten, M.S., Krijger, P.H.L., Teunissen, H., Medema, R.H., van Steensel, B., Brummelkamp, T.R., de Wit, E., Rowland, B.D., 2017. The Cohesin Release Factor WAPL Restricts Chromatin Loop Extension. Cell 169, 693–707.e14. 10.1016/j.cell.2017.04.013

Hajjoul, H., Mathon, J., Ranchon, H., Goiffon, I., Mozziconacci, J., Albert, B., Carrivain, P., Victor, J.-M., Gadal, O., Bystricky, K., Bancaud, A., 2013. High-throughput chromatin motion tracking in living yeast reveals the flexibility of the fiber throughout the genome. Genome Res. 23, 1829–1838. 10.1101/gr.157008.113

Hansen, A.S., Pustova, I., Cattoglio, C., Tjian, R., Darzacq, X., 2017. CTCF and cohesin regulate chromatin loop stability with distinct dynamics. eLife 6, e25776. 10.7554/eLife.25776

Hill, L., Ebert, A., Jaritz, M., Wutz, G., Nagasaka, K., Tagoh, H., Kostanova-Poliakova, D., Schindler, K., Sun, Q., Bönelt, P., Fischer, M., Peters, J.-M., Busslinger, M., 2020. Wapl repression by Pax5 promotes V gene recombination by Igh loop extrusion. Nature 584, 142–147. 10.1038/s41586-020-2454-y

Holzmann, J., Politi, A.Z., Nagasaka, K., Hantsche-Grininger, M., Walther, N., Koch, B., Fuchs, J., Dürnberger, G., Tang, W., Ladurner, R., Stocsits, R.R., Busslinger, G.A., Novák, B., Mechtler, K., Davidson, I.F., Ellenberg, J., Peters, J.-M., 2019. Absolute quantification of cohesin, CTCF and their regulators in human cells. eLife 8, e46269. 10.7554/eLife.46269

Huis in ’t Veld, P.J., Herzog, F., Ladurner, R., Davidson, I.F., Piric, S., Kreidl, E., Bhaskara, V., Aebersold, R., Peters, J.-M., 2014. Characterization of a DNA exit gate in the human cohesin ring. Science 346, 968–972. 10.1126/science.1256904

James, M.B., Giorgio, T.D., 2000. Nuclear-associated plasmid, but not cell-associated plasmid, is correlated with transgene expression in cultured mammalian cells. Mol. Ther. J. Am. Soc. Gene Ther. 1, 339–346. 10.1006/mthe.2000.0054

Kesel, A.J., Day, C.W., Montero, C.M., Schinazi, R.F., 2016. A new oxygen modification cyclooctaoxygen binds to nucleic acids as sodium crown complex. Biochim. Biophys. Acta 1860, 785–794. 10.1016/j.bbagen.2016.01.022

Kiefer, L., Chiosso, A., Langen, J., Buckley, A., Gaudin, S., Rajkumar, S.M., Servito, G.I.F., Cha, E.S., Vijay, A., Yeung, A., Horta, A., Mui, M.H., Canzio, D., 2023. WAPL functions as a rheostat of Protocadherin isoform diversity that controls neural wiring. Science 380, eadf8440. 10.1126/science.adf8440

Kikuchi, S., Borek, D.M., Otwinowski, Z., Tomchick, D.R., Yu, H., 2016. Crystal structure of the cohesin loader Scc2 and insight into cohesinopathy. Proc. Natl. Acad. Sci. U. S. A. 113, 12444–12449. 10.1073/pnas.1611333113

Kim, Y., Shi, Z., Zhang, H., Finkelstein, I.J., Yu, H., 2019. Human cohesin compacts DNA by loop extrusion. Science 366, 1345–1349. 10.1126/science.aaz4475

Kueng, S., Hegemann, B., Peters, B.H., Lipp, J.J., Schleiffer, A., Mechtler, K., Peters, J.-M., 2006. Wapl controls the dynamic association of cohesin with chromatin. Cell 127, 955–967. 10.1016/j.cell.2006.09.040

Ladurner, R., Kreidl, E., Ivanov, M.P., Ekker, H., Idarraga-Amado, M.H., Busslinger, G.A., Wutz, G., Cisneros, D.A., Peters, J.-M., 2016. Sororin actively maintains sister chromatid cohesion. EMBO J. 35, 635–653. 10.15252/embj.201592532

Losada, A., Yokochi, T., Hirano, T., 2005. Functional contribution of Pds5 to cohesin-mediated cohesion in human cells and Xenopus egg extracts. J. Cell Sci. 118, 2133–2141. 10.1242/jcs.02355

Luppino, J.M., Field, A., Nguyen, S.C., Park, D.S., Shah, P.P., Abdill, R.J., Lan, Y., Yunker, R., Jain, R., Adelman, K., Joyce, E.F., 2022. Co-depletion of NIPBL and WAPL balance cohesin activity to correct gene misexpression. PLoS Genet. 18, e1010528. 10.1371/journal.pgen.1010528

Mach, P., Kos, P.I., Zhan, Y., Cramard, J., Gaudin, S., Tünnermann, J., Marchi, E., Eglinger, J., Zuin, J., Kryzhanovska, M., Smallwood, S., Gelman, L., Roth, G., Nora, E.P., Tiana, G., Giorgetti, L., 2022. Cohesin and CTCF control the dynamics of chromosome folding. Nat. Genet. 54, 1907–1918. 10.1038/s41588-022-01232-7

McCord, R.P., Kaplan, N., Giorgetti, L., 2020. Chromosome Conformation Capture and Beyond: Toward an Integrative View of Chromosome Structure and Function. Mol. Cell 77, 688–708. 10.1016/j.molcel.2019.12.021

Muir, K.W., Kschonsak, M., Li, Y., Metz, J., Haering, C.H., Panne, D., 2016. Structure of the Pds5-Scc1 Complex and Implications for Cohesin Function. Cell Rep. 14, 2116–2126. 10.1016/j.celrep.2016.01.078

Murayama, Y., Uhlmann, F., 2014. Biochemical reconstitution of topological DNA binding by the cohesin ring. Nature 505, 367–371. 10.1038/nature12867

Nasmyth, K., 2011. Cohesin: a catenase with separate entry and exit gates? Nat. Cell Biol. 13, 1170–1177. 10.1038/ncb2349

Nasmyth, K.A., Lee, B.-G., Roig, M.B., Löwe, J., 2023. What AlphaFold tells us about cohesin’s retention on and release from chromosomes. eLife 12, RP88656. 10.7554/eLife.88656

Naumova, N., Imakaev, M., Fudenberg, G., Zhan, Y., Lajoie, B.R., Mirny, L.A., Dekker, J., 2013. Organization of the mitotic chromosome. Science 342, 948–953. 10.1126/science.1236083

Nomidis, S.K., Carlon, E., Gruber, S., Marko, J.F., 2022. DNA tension-modulated translocation and loop extrusion by SMC complexes revealed by molecular dynamics simulations. Nucleic Acids Res. 50, 4974–4987. 10.1093/nar/gkac268

Nuebler, J., Fudenberg, G., Imakaev, M., Abdennur, N., Mirny, L.A., 2018. Chromatin organization by an interplay of loop extrusion and compartmental segregation. Proc. Natl. Acad. Sci. U. S. A. 115, E6697–E6706. 10.1073/pnas.1717730115

Onn, I., Heidinger-Pauli, J.M., Guacci, V., Unal, E., Koshland, D.E., 2008. Sister chromatid cohesion: a simple concept with a complex reality. Annu. Rev. Cell Dev. Biol. 24, 105–129. 10.1146/annurev.cellbio.24.110707.175350

Open2C, Abdennur, N., Fudenberg, G., Flyamer, I.M., Galitsyna, A.A., Goloborodko, A., Imakaev, M., Venev, S.V., 2024. Pairtools: From sequencing data to chromosome contacts. PLoS Comput. Biol. 20, e1012164. 10.1371/journal.pcbi.1012164

Ouyang, Z., Yu, H., 2017. Releasing the cohesin ring: A rigid scaffold model for opening the DNA exit gate by Pds5 and Wapl. BioEssays News Rev. Mol. Cell. Dev. Biol. 39. 10.1002/bies.201600207

Ouyang, Z., Zheng, G., Song, J., Borek, D.M., Otwinowski, Z., Brautigam, C.A., Tomchick, D.R., Rankin, S., Yu, H., 2013. Structure of the human cohesin inhibitor Wapl. Proc. Natl. Acad. Sci. U. S. A. 110, 11355–11360. 10.1073/pnas.1304594110

Panizza, S., Tanaka, T., Hochwagen, A., Eisenhaber, F., Nasmyth, K., 2000. Pds5 cooperates with cohesin in maintaining sister chromatid cohesion. Curr. Biol. CB 10, 1557–1564. 10.1016/s0960-9822(00)00854-x

Petela, N.J., Gligoris, T.G., Metson, J., Lee, B.-G., Voulgaris, M., Hu, B., Kikuchi, S., Chapard, C., Chen, W., Rajendra, E., Srinivisan, M., Yu, H., Löwe, J., Nasmyth, K.A., 2018. Scc2 Is a Potent Activator of Cohesin’s ATPase that Promotes Loading by Binding Scc1 without Pds5. Mol. Cell 70, 1134–1148.e7. 10.1016/j.molcel.2018.05.022

Peters, J.-M., Nishiyama, T., 2012. Sister Chromatid Cohesion. Cold Spring Harb. Perspect. Biol. 4, a011130. 10.1101/cshperspect.a011130

Pezic, D., Weeks, S.L., Hadjur, S., 2017. More to cohesin than meets the eye: complex diversity for fine-tuning of function. Curr. Opin. Genet. Dev. 43, 93–100. 10.1016/j.gde.2017.01.004

Phillips, C.L., Anderson, J.A., Glotzer, S.C., 2011. Pseudo-random number generation for Brownian Dynamics and Dissipative Particle Dynamics simulations on GPU devices. J. Comput. Phys. 230, 7191–7201. 10.1016/j.jcp.2011.05.021

Pobegalov, G., Chu, L.-Y., Peters, J.-M., Molodtsov, M.I., 2023. Single cohesin molecules generate force by two distinct mechanisms. Nat. Commun. 14, 3946. 10.1038/s41467-023-39696-8

Polovnikov, K.E., Slavov, B., Belan, S., Imakaev, M., Brandão, H.B., Mirny, L.A., 2023. Crumpled polymer with loops recapitulates key features of chromosome organization. Phys. Rev. X 13, 041029. 10.1103/physrevx.13.041029

Rhodes, J., Mazza, D., Nasmyth, K., Uphoff, S., 2017. Scc2/Nipbl hops between chromosomal cohesin rings after loading. eLife 6, e30000. 10.7554/eLife.30000

Samejima, K., Gibcus, J.H., Abraham, S., Cisneros-Soberanis, F., Samejima, I., Beckett, A.J., Pučeková, N., Abad, M.A., Medina-Pritchard, B., Paulson, J.R., Xie, L., Jeyaprakash, A.A., Prior, I.A., Mirny, L.A., Dekker, J., Goloborodko, A., Earnshaw, W.C., 2024. Rules of engagement for condensins and cohesins guide mitotic chromosome formation. bioRxiv 2024.04.18.590027. 10.1101/2024.04.18.590027

Shintomi, K., Hirano, T., 2009. Releasing cohesin from chromosome arms in early mitosis: opposing actions of Wapl-Pds5 and Sgo1. Genes Dev. 23, 2224–2236. 10.1101/gad.1844309

Srinivasan, M., Petela, N.J., Scheinost, J.C., Collier, J., Voulgaris, M., B Roig, M., Beckouët, F., Hu, B., Nasmyth, K.A., 2019. Scc2 counteracts a Wapl-independent mechanism that releases cohesin from chromosomes during G1. eLife 8, e44736. 10.7554/eLife.44736

Sun, Yuao, Xu, X., Zhao, W., Zhang, Y., Chen, K., Li, Y., Wang, X., Zhang, M., Xue, B., Yu, W., Hou, Y., Wang, C., Xie, W., Li, C., Kong, D., Wang, S., Sun, Yujie, 2023. RAD21 is the core subunit of the cohesin complex involved in directing genome organization. Genome Biol. 24, 155. 10.1186/s13059-023-02982-1

Tedeschi, A., Wutz, G., Huet, S., Jaritz, M., Wuensche, A., Schirghuber, E., Davidson, I.F., Tang, W., Cisneros, D.A., Bhaskara, V., Nishiyama, T., Vaziri, A., Wutz, A., Ellenberg, J., Peters, J.-M., 2013. Wapl is an essential regulator of chromatin structure and chromosome segregation. Nature 501, 564–568. 10.1038/nature12471

van Ruiten, M.S., Rowland, B.D., 2021. On the choreography of genome folding: A grand pas de deux of cohesin and CTCF. Curr. Opin. Cell Biol. 70, 84–90. 10.1016/j.ceb.2020.12.001

van Ruiten, M.S., van Gent, D., Sedeño Cacciatore, Á., Fauster, A., Willems, L., Hekkelman, M.L., Hoekman, L., Altelaar, M., Haarhuis, J.H.I., Brummelkamp, T.R., de Wit, E., Rowland, B.D., 2022. The cohesin acetylation cycle controls chromatin loop length through a PDS5A brake mechanism. Nat. Struct. Mol. Biol. 29, 586–591. 10.1038/s41594-022-00773-z

Wutz, G., Ladurner, R., St Hilaire, B.G., Stocsits, R.R., Nagasaka, K., Pignard, B., Sanborn, A., Tang, W., Várnai, C., Ivanov, M.P., Schoenfelder, S., van der Lelij, P., Huang, X., Dürnberger, G., Roitinger, E., Mechtler, K., Davidson, I.F., Fraser, P., Lieberman-Aiden, E., Peters, J.-M., 2020. ESCO1 and CTCF enable formation of long chromatin loops by protecting cohesinSTAG1 from WAPL. eLife 9, e52091. 10.7554/eLife.52091

Wutz, G., Várnai, C., Nagasaka, K., Cisneros, D.A., Stocsits, R.R., Tang, W., Schoenfelder, S., Jessberger, G., Muhar, M., Hossain, M.J., Walther, N., Koch, B., Kueblbeck, M., Ellenberg, J., Zuber, J., Fraser, P., Peters, J.-M., 2017. Topologically associating domains and chromatin loops depend on cohesin and are regulated by CTCF, WAPL, and PDS5 proteins. EMBO J. 36, 3573–3599. 10.15252/embj.201798004

Zhang, B., Chang, J., Fu, M., Huang, J., Kashyap, R., Salavaggione, E., Jain, S., Kulkarni, S., Deardorff, M.A., Uzielli, M.L.G., Dorsett, D., Beebe, D.C., Jay, P.Y., Heuckeroth, R.O., Krantz, I., Milbrandt, J., 2009. Dosage effects of cohesin regulatory factor PDS5 on mammalian development: implications for cohesinopathies. PloS One 4, e5232. 10.1371/journal.pone.0005232

Zhang, N., Coutinho, L.E., Pati, D., 2021. PDS5A and PDS5B in Cohesin Function and Human Disease. Int. J. Mol. Sci. 22, 5868. 10.3390/ijms22115868

