## Supplemental Information for "The physical chemistry of interphase loop extrusion"

(Dated: August 23, 2024)

### I. RATE MAPPING PROCEDURE

Using the notations of the main text, the chemical reaction network outlined in Fig. 1c explicitly reads as:

$$\begin{aligned}
\mathrm{R}_{\mathrm{free}} + \mathrm{N}_{\mathrm{free}} &\xrightarrow{k_{\mathrm{on}}} \mathrm{RN}, \\
\mathrm{RN} &\underset{k_{\mathrm{RN}}}{\overset{k_{\mathrm{NR}}}{\rightleftharpoons}} \mathrm{R} + \mathrm{N}_{\mathrm{free}}, \\
\mathrm{R} + \mathrm{P}_{\mathrm{free}} &\underset{k_{\mathrm{PR}}}{\overset{k_{\mathrm{RP}}}{\rightleftharpoons}} \mathrm{RP}, \\
\mathrm{RP} + \mathrm{W}_{\mathrm{free}} &\underset{k_{\mathrm{WP}}}{\overset{k_{\mathrm{PW}}}{\rightleftharpoons}} \mathrm{RW} + \mathrm{P}_{\mathrm{free}}, \\
\mathrm{RW} &\xrightarrow{k_{\mathrm{off}}} \mathrm{R}_{\mathrm{free}} + \mathrm{W}_{\mathrm{free}},
\end{aligned}$$

where the "free" subscript denotes species not bound to chromatin. In the framework of the law of mass action, the kinetic equations describing the time evolution of the free concentration of each cohesin subunit thus take the form of a system of coupled ODEs,

$$\frac{\partial [\mathrm{R}]_{\mathrm{free}}}{\partial t} = k_{\mathrm{off}} [\mathrm{RW}] - k_{\mathrm{on}} [\mathrm{R}]_{\mathrm{free}} [\mathrm{N}]_{\mathrm{free}}, \tag{1}$$

$$\frac{\partial [\mathrm{N}]_{\mathrm{free}}}{\partial t} = k_{\mathrm{NR}} [\mathrm{RN}] - k_{\mathrm{RN}} [\mathrm{R}] [\mathrm{N}]_{\mathrm{free}} - k_{\mathrm{on}} [\mathrm{R}]_{\mathrm{free}} [\mathrm{N}]_{\mathrm{free}}, \tag{2}$$

$$\frac{\partial [\mathrm{P}]_{\mathrm{free}}}{\partial t} = k_{\mathrm{PR}} [\mathrm{RP}] + k_{\mathrm{PW}} [\mathrm{RP}] [\mathrm{W}]_{\mathrm{free}} - k_{\mathrm{RP}} [\mathrm{R}] [\mathrm{P}]_{\mathrm{free}} - k_{\mathrm{WP}} [\mathrm{RW}] [\mathrm{P}]_{\mathrm{free}}, \tag{3}$$

$$\frac{\partial [\mathrm{W}]_{\mathrm{free}}}{\partial t} = k_{\mathrm{off}} [\mathrm{RW}] + k_{\mathrm{WP}} [\mathrm{RW}] [\mathrm{P}]_{\mathrm{free}} - k_{\mathrm{PW}} [\mathrm{RP}] [\mathrm{W}]_{\mathrm{free}}. \tag{4}$$

Let us consider the chromatin association/dissociation reaction of a generic species X,

$$\mathrm{X}_{\mathrm{free}} \underset{k_{\mathrm{off}}^{\mathrm{X}}}{\overset{k_{\mathrm{on}}^{\mathrm{X}}}{\rightleftharpoons}} \mathrm{X}_{\mathrm{bound}}.$$

In the limit of large excess of chromatin binding sites, the corresponding kinetic equation for the free X population reads as

$$\frac{\partial [\mathrm{X}]_{\mathrm{free}}}{\partial t} = k_{\mathrm{off}}^{\mathrm{X}} [\mathrm{X}]_{\mathrm{bound}} - k_{\mathrm{on}}^{\mathrm{X}} [\mathrm{X}]_{\mathrm{free}}. \tag{5}$$

Let us denote by $[\mathrm{X}]_{\mathrm{tot}} \equiv [\mathrm{X}]_{\mathrm{bound}} + [\mathrm{X}]_{\mathrm{free}}$ the total nuclear content of X. Assuming $[\mathrm{X}]_{\mathrm{tot}}$ to be constant throughout the G1 stage of the cell cycle, to which we restrict our current study, the equilibrium bound fraction $f_{\mathrm{X}}$ may be obtained by solving Eq. (5) at steady state,

$$f_{\mathrm{X}} \equiv \frac{\left[\mathrm{X}\right]_{\text{bound}}^{\text{eq}}}{\left[\mathrm{X}\right]_{\text{tot}}} = 1 - \frac{\left[\mathrm{X}\right]_{\text{free}}^{\text{eq}}}{\left[\mathrm{X}\right]_{\text{tot}}} = \frac{k_{\text{on}}^{\mathrm{X}}}{k_{\text{on}}^{\mathrm{X}} + k_{\text{off}}^{\mathrm{X}}}, \tag{6}$$

while the unbinding rate $k_{\text{off}}^{\mathrm{X}}$ is related to the chromatin residence time $\tau_{\mathrm{X}}$ via

$$\tau_{\mathrm{X}} = \frac{1}{k_{\text{off}}^{\mathrm{X}}}. \tag{7}$$

Substituting for species X the cohesin subunits RAD21, NIPBL, PDS5 & WAPL, a direct term-by-term comparison of Eqs. (1)–(4) with Eq. (5) yields

$$k_{\text{on}}^{\mathrm{R}} = k_{\text{on}}\left[\mathrm{N}\right]_{\text{free}}, \tag{8}$$

$$k_{\text{off}}^{\mathrm{R}} = k_{\text{off}}\frac{\left[\mathrm{RW}\right]}{\left[\mathrm{R}\right]_{\text{bound}}}, \tag{9}$$

$$k_{\text{on}}^{\mathrm{N}} = k_{\mathrm{RN}}\left[\mathrm{R}\right] + k_{\text{on}}\left[\mathrm{R}\right]_{\text{free}}, \tag{10}$$

$$k_{\text{off}}^{\mathrm{N}} = k_{\mathrm{NR}}, \tag{11}$$

$$k_{\text{on}}^{\mathrm{P}} = k_{\mathrm{RP}}\left[\mathrm{R}\right] + k_{\mathrm{WP}}\left[\mathrm{RW}\right], \tag{12}$$

$$k_{\text{off}}^{\mathrm{P}} = k_{\mathrm{PR}} + k_{\mathrm{PW}}\left[\mathrm{W}\right]_{\text{free}}, \tag{13}$$

$$k_{\text{on}}^{\mathrm{W}} = k_{\mathrm{PW}}\left[\mathrm{RP}\right], \tag{14}$$

$$k_{\text{off}}^{\mathrm{W}} = k_{\text{off}} + k_{\mathrm{WP}}\left[\mathrm{P}\right]_{\text{free}}, \tag{15}$$

where we used $\left[\mathrm{N}\right]_{\text{bound}} = \left[\mathrm{RN}\right]$, $\left[\mathrm{P}\right]_{\text{bound}} = \left[\mathrm{RP}\right]$ and $\left[\mathrm{W}\right]_{\text{bound}} = \left[\mathrm{RW}\right]$. Plugging in Eqs. (6) and (7), Eqs. (8)–(15) may be recast in the form, at chemical equilibrium,

$$\frac{1}{\tau_{\mathrm{R}}}\frac{f_{\mathrm{R}}}{1-f_{\mathrm{R}}} = k_{\text{on}}\left(1-f_{\mathrm{N}}\right)\left[\mathrm{N}\right]_{\text{tot}}, \tag{16}$$

$$\frac{1}{\tau_{\mathrm{R}}} = k_{\text{off}}\frac{f_{\mathrm{W}}\left[\mathrm{W}\right]_{\text{tot}}}{f_{\mathrm{R}}\left[\mathrm{R}\right]_{\text{tot}}}, \tag{17}$$

$$\frac{1}{\tau_{\mathrm{N}}}\frac{f_{\mathrm{N}}}{1-f_{\mathrm{N}}} = k_{\mathrm{RN}}\Big(f_{\mathrm{R}}\left[\mathrm{R}\right]_{\text{tot}} - f_{\mathrm{N}}\left[\mathrm{N}\right]_{\text{tot}} - f_{\mathrm{P}}\left[\mathrm{P}\right]_{\text{tot}} - f_{\mathrm{W}}\left[\mathrm{W}\right]_{\text{tot}}\Big) + k_{\text{on}}\left(1-f_{\mathrm{R}}\right)\left[\mathrm{R}\right]_{\text{tot}}, \tag{18}$$

$$\frac{1}{\tau_{\mathrm{N}}} = k_{\mathrm{NR}}, \tag{19}$$

$$\frac{1}{\tau_{\mathrm{P}}}\frac{f_{\mathrm{P}}}{1-f_{\mathrm{P}}} = k_{\mathrm{RP}}\Big(f_{\mathrm{R}}\left[\mathrm{R}\right]_{\text{tot}} - f_{\mathrm{N}}\left[\mathrm{N}\right]_{\text{tot}} - f_{\mathrm{P}}\left[\mathrm{P}\right]_{\text{tot}} - f_{\mathrm{W}}\left[\mathrm{W}\right]_{\text{tot}}\Big) + k_{\mathrm{WP}}f_{\mathrm{W}}\left[\mathrm{W}\right]_{\text{tot}}, \tag{20}$$

$$\frac{1}{\tau_{\mathrm{P}}} = k_{\mathrm{PR}} + k_{\mathrm{PW}}\left(1-f_{\mathrm{W}}\right)\left[\mathrm{W}\right]_{\text{tot}}, \tag{21}$$

$$\frac{1}{\tau_{\mathrm{W}}}\frac{f_{\mathrm{W}}}{1-f_{\mathrm{W}}} = k_{\mathrm{PW}}f_{\mathrm{P}}\left[\mathrm{P}\right]_{\text{tot}}, \tag{22}$$

$$\frac{1}{\tau_{\mathrm{W}}} = k_{\text{off}} + k_{\mathrm{WP}}\left(1-f_{\mathrm{P}}\right)\left[\mathrm{P}\right]_{\text{tot}}, \tag{23}$$

TABLE S1. **Equilibrium rates of the five-state model for wild-type HeLa cells** (c.f. Fig. 1 and Table 1 of the main text).

| State transition rates | |
|---|---|
| $k_{\mathrm{on}}$ | $3.4 \times 10^{-5}\,\mathrm{s}^{-1}\,\mathrm{mol}^{-1}$ |
| $k_{\mathrm{off}}$ | $9.2 \times 10^{-3}\,\mathrm{s}^{-1}$ |
| $k_{\mathrm{NR}}$ | $1.4 \times 10^{-2}\,\mathrm{s}^{-1}$ |
| $k_{\mathrm{RN}}$ | $2.0 \times 10^{-4}\,\mathrm{s}^{-1}\,\mathrm{mol}^{-1}$ |
| $k_{\mathrm{RP}}$ | $2.7 \times 10^{-4}\,\mathrm{s}^{-1}\,\mathrm{mol}^{-1}$ |
| $k_{\mathrm{PR}}$ | $7.4 \times 10^{-3}\,\mathrm{s}^{-1}$ |
| $k_{\mathrm{PW}}$ | $1.6 \times 10^{-4}\,\mathrm{s}^{-1}\,\mathrm{mol}^{-1}$ |
| $k_{\mathrm{WP}}$ | $1.4 \times 10^{-4}\,\mathrm{s}^{-1}\,\mathrm{mol}^{-1}$ |

in which we substituted $\left[\mathrm{R}\right] = f_{\mathrm{R}}\left[\mathrm{R}\right]_{\mathrm{tot}} - f_{\mathrm{N}}\left[\mathrm{N}\right]_{\mathrm{tot}} - f_{\mathrm{P}}\left[\mathrm{P}\right]_{\mathrm{tot}} - f_{\mathrm{W}}\left[\mathrm{W}\right]_{\mathrm{tot}}$ based on the mass conservation of RAD21.

Using the experimentally-determined bound fractions $f_{\mathrm{X}}$ and residence times $\tau_{\mathrm{X}}$ estimated from FRAP for each of the 4 relevant cohesin subunits, as well as the absolute G1 protein numbers $\left[\mathrm{X}\right]_{\mathrm{tot}}$ obtained as described in the main text, Eqs. (16)–(23) yield a linear system of 8 coupled equations involving the 8 unknown rates $k_{\mathrm{on}}$, $k_{\mathrm{off}}$, $k_{\mathrm{NR}}$, $k_{\mathrm{RN}}$, $k_{\mathrm{RP}}$, $k_{\mathrm{PR}}$, $k_{\mathrm{PW}}$, $k_{\mathrm{WP}}$ governing the chemical reaction network (Fig. 1c). Eqs. (16)–(23) were inverted symbolically using the *SymPy* library, and the computed values for the rate $k$ were plugged into Eqs. (1)–(4), which were then integrated numerically as described in the main text. For the combinatorial exploration of reaction networks, all possible permutations of bound cohesin state sequences were systematically generated using the *itertools* library, and the corresponding kinetic (Eqs. (1)–(4)) and rate-mapping (Eqs. (16)–(23)) equations were derived programmatically and similarly solved using *SymPy*.

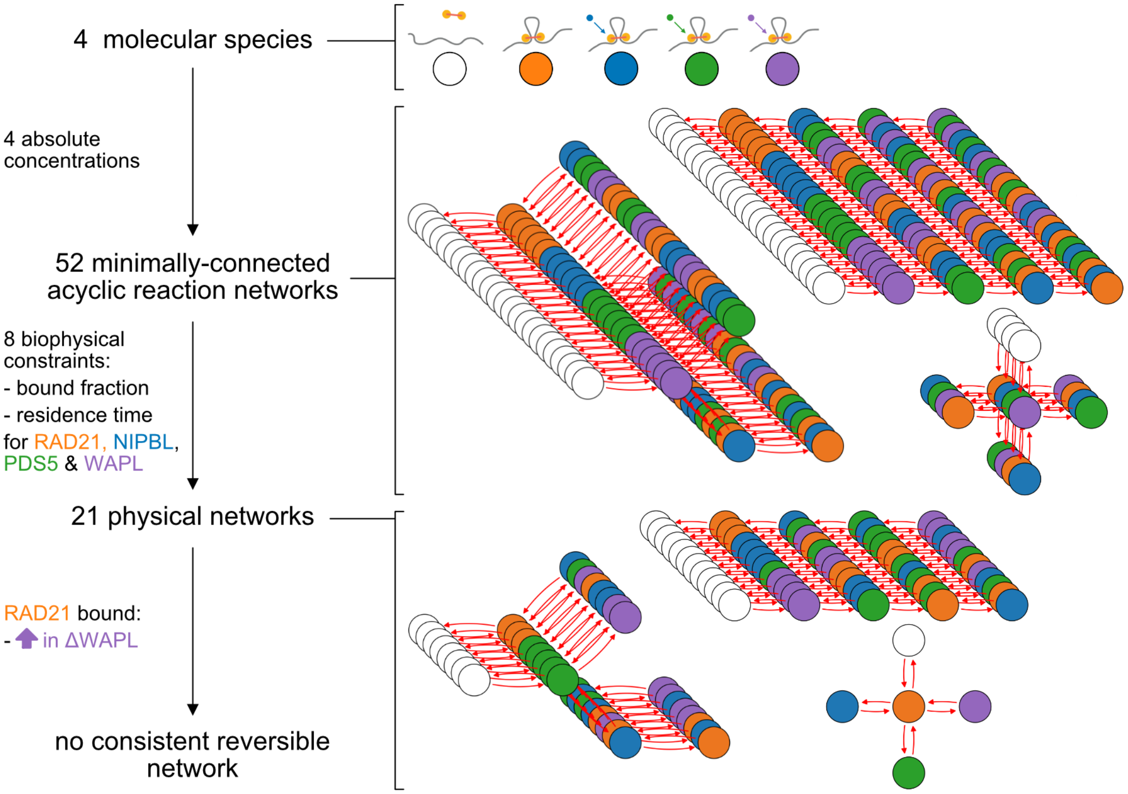


FIG. S1. **Chromatin entry and exit involves distinct cohesin molecular pathways.** To systematically explore and rule out the possibility of cohesin loading and unloading via a single pathway, we apply the same decimation procedure as used in Fig. 1 of the main text to all possible acyclic, fully reversible networks with minimal number of edges (8). While 21 networks can be found with non-negative rates, none of these networks lead to an increase in the bound fraction of RAD21 upon depletion of WAPL — or, more generally, of any of the other cohesin regulators. Thus, no fully reversible networks are consistent with experimental observations

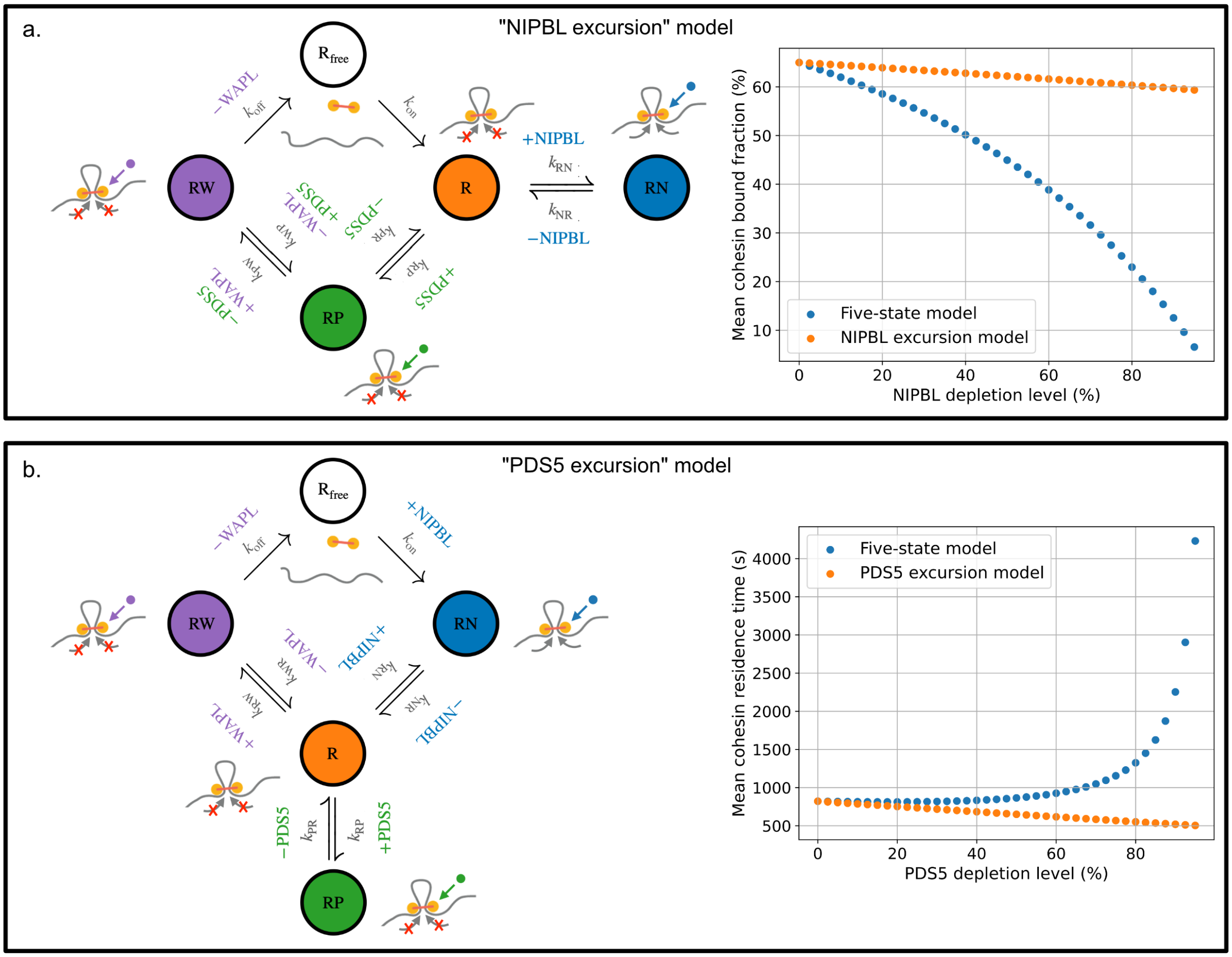


FIG. S2. **Alternate topologies are inconsistent with experiments.**

**a.** NIPBL excursion model, where NIPBL is not involved with loading but instead binds reversibly after the core complex has bound chromatin. For this topology, NIPBL depletion does not substantially lower the bound fraction, unlike experimental observations. The inconsistency of this alternate topology provides mathematical support for the role of NIPBL in productive cohesin loading in an extrusion cycle. **b.** PDS5 excursion model, where PDS5 reversibly binds the core complex and does not promote WAPL binding. For this topology, PDS5 depletion actually slightly lowers RAD21 residence time, instead of increasing RAD21 residence time as observed experimentally. The inconsistency of this alternate topology argues that PDS5 is positioned along the reaction cycle in such a way to influence unloading rates.

| | | | Mutant depletion | | |
|---|---|---|---|---|---|
| | | | ΔNIPBL | ΔPDS5 | ΔWAPL |
| Cohesin accessory protein | RAD21 | Bound fraction | *In vivo* [1] | *In vivo* [1] | *In vivo* [1] |
| | | Residence time | *In vitro* [2] | *In vivo* [1] | *In vivo* [1] |
| | NIPBL | Bound fraction | | | *In vivo* [3] |
| | | Residence time | *In vitro* [2] | | *In vivo* [3] |
| | PDS5 | Bound fraction | | | |
| | | Residence time | | | |
| | WAPL | Bound fraction | | | |
| | | Residence time | | | |

TABLE S2. **Predicted impacts of accessory protein depletions on their chromatin association dynamics in the five-state model.**

Recapitulative table of the role of various cohesin accessory protein depletions (columns) on the chromatin-associated fraction and residence time of other proteins (rows). Red (resp. blue) colors signify that the model predicts an increase (resp. decrease) of the corresponding quantity in the different mutants relative to its magnitude in wild-type HeLa cells. Gray shades mark a lack of significant deviation from the wild-type value. References point to experimental studies reporting *in vivo* or *in vitro* validation of the predicted changes, wherever available. We note that NIPBL depletion leads to a drastic reduction in the bound fractions of PDS5 and WAPL, consistent with a significant inhibition of cohesin loading, but is associated with a more moderate drop (10-20%) in their respective cohesin residence times (Fig. 3c of the main text).

| Cohesin accessory protein | | Mutant depletion | | |
|---|---|---|---|---|
| | | ΔNIPBL | ΔPDS5 | ΔWAPL |
| RAD21 | Bound fraction | *In vivo* [1] | *In vivo* [1] | *In vivo* [1] |
| RAD21 | Residence time | *In vitro* [2] | *In vivo* [1] | *In vivo* [1] |
| NIPBL | Bound fraction | | | *In vivo* [3] |
| NIPBL | Residence time | *In vitro* [2] | | *In vivo* [3] |
| PDS5 | Bound fraction | | | |
| PDS5 | Residence time | | | |
| WAPL | Bound fraction | | | |
| WAPL | Residence time | | | |

TABLE S3. **Same as Table S2 for the PDS5-WAPL co-bound model** (Fig. S5a). Note that the main qualitative difference with the five-state model lies in the residence time of PDS5 in WAPL-depleted cells, which decreases relative to wild-type in the PDS5-WAPL co-bound model, but increases in the case of the strict subunit exchange assumed by the five-state model (Fig. S5d).

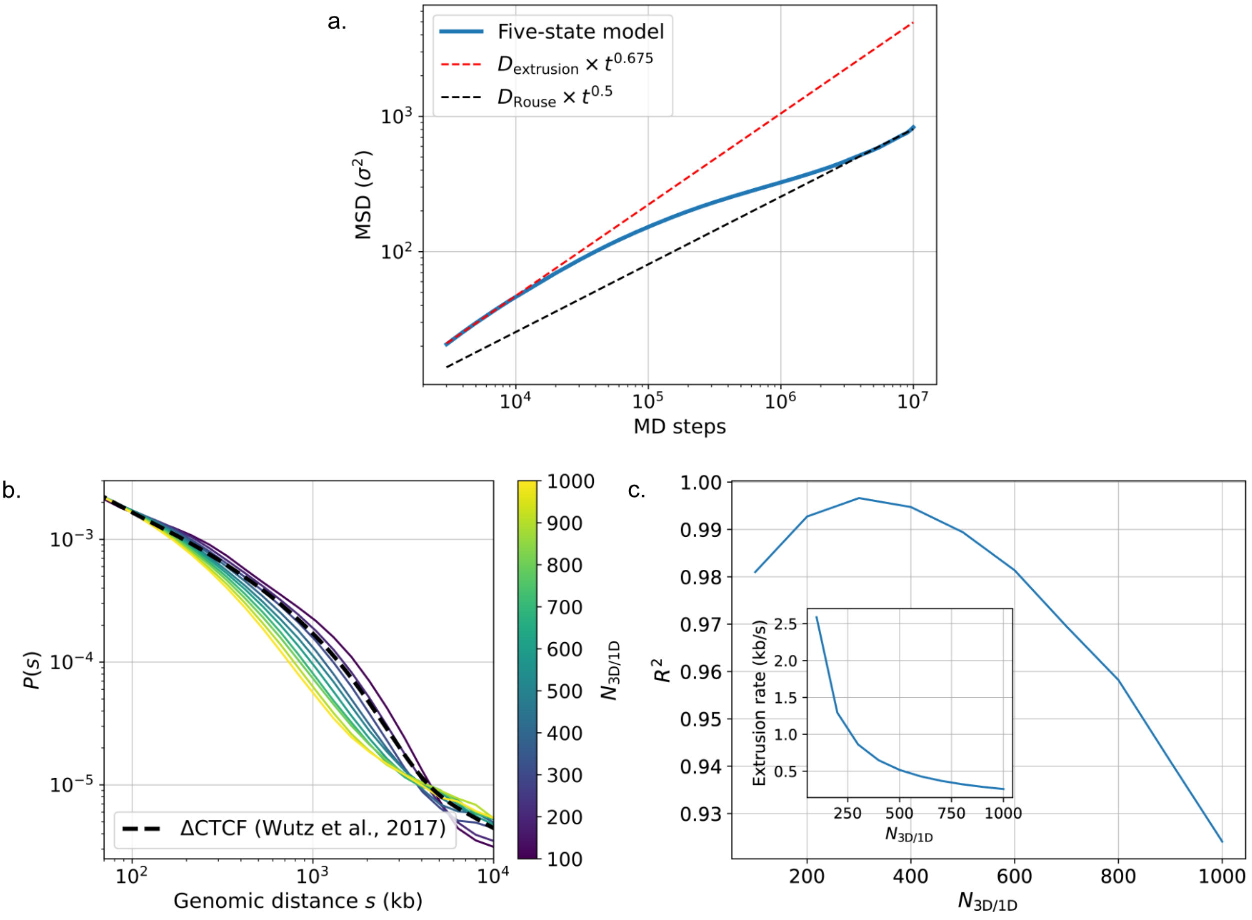


FIG. S3. **Time mapping & influence of loop extrusion rate.**

**a**: Mean-squared displacement (MSD) of individual monomers as predicted by the five-state model at wild-type HeLa protein expression levels. The correspondence between model and experimental time units (1 MD step $\sim$ 5 ms) is obtained by comparing the slopes of the long- and short-time asymptotes to the respective experimental values $D_{\text{Rouse}} \simeq 0.01\,\mu\text{m}^2/\text{s}^{0.5}$ and $D_{\text{extrusion}} \simeq 0.0075\,\mu\text{m}^2/\text{s}^{0.675}$, as estimated in budding yeast and CTCF-depleted mESCs. **b**: Computational contact-vs-distance curves ($P(s)$) at different ratios of 3D-to-1D steps ($N_{3D/1D}$). Dashed line: experimental profile obtained in ΔCTCF mutants [1]. **c**: Mean-squared $R^2$ coefficient in the model vs. experimental P(s) curves, averaged over the distance range [50 kb: 5,000 kb]. Inset: Numerical correspondence between $N_{3D/1D}$ and mean extrusion rate ($v$) in wild-type HeLa cells.

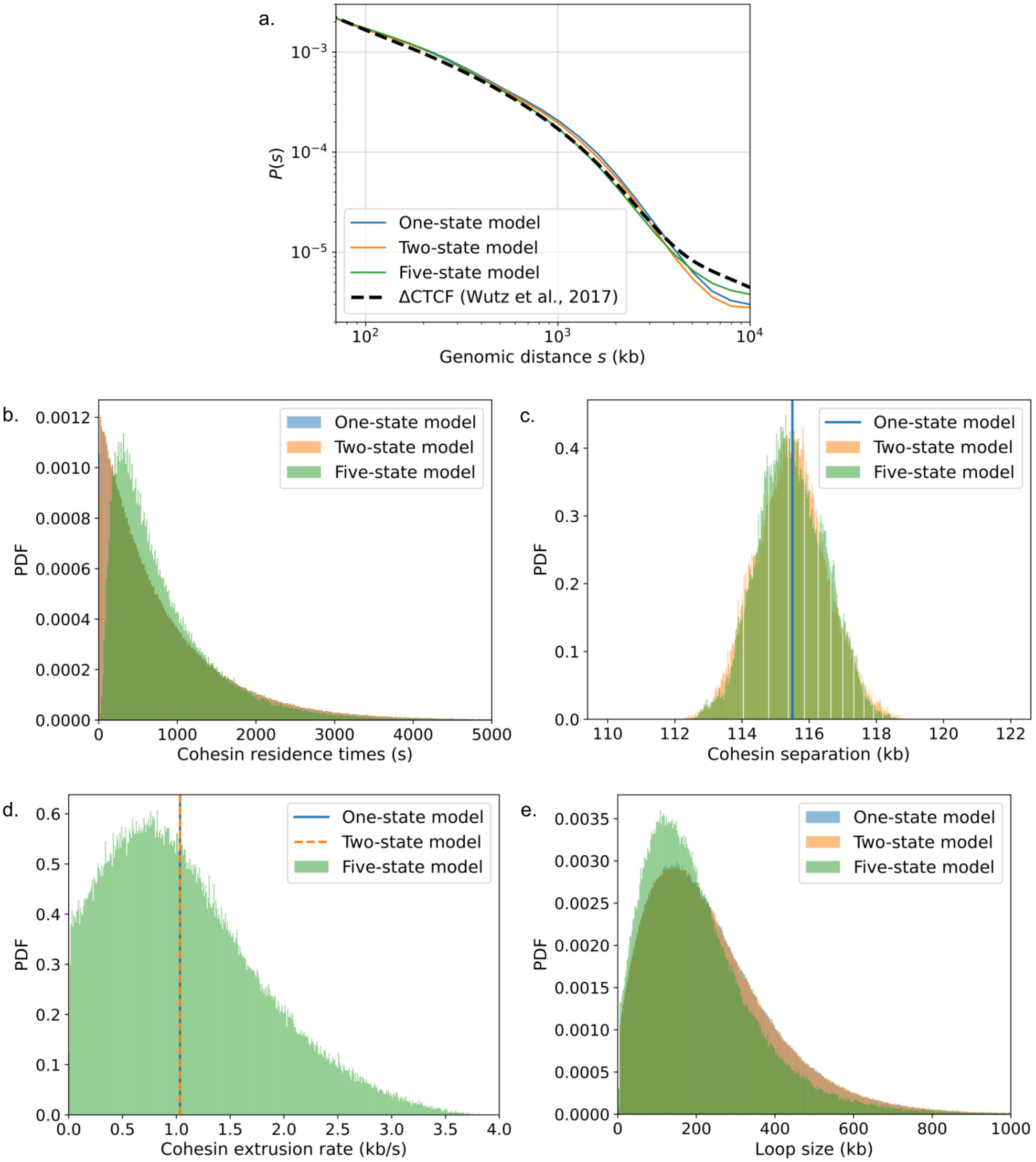
a.
$10^{-3}$
$10^{-4}$
$10^{-5}$
$P(s)$
One-state model
Two-state model
Five-state model
ΔCTCF (Wutz et al., 2017)
$10^2$
$10^3$
$10^4$
Genomic distance $s$ (kb)
b.
PDF
One-state model
Two-state model
Five-state model
Cohesin residence times (s)
c.
PDF
One-state model
Two-state model
Five-state model
Cohesin separation (kb)
d.
PDF
One-state model
Two-state model
Five-state model
Cohesin extrusion rate (kb/s)
e.
PDF
One-state model
Two-state model
Five-state model
Loop size (kb)

FIG. S4. **The five-state model provides additional sources of heterogeneity absent from previous cohesin models.**

One state and two-state models are parameterized to have the same average extrusion rate, cohesin residence time and number of loaded extruders as the five-state model, but respectively assume a constant extrusion rate with or without immediate cohesin rebinding. **a.** Contact frequency versus distance curves are largely similar across models. **b.** The distribution of residence times of the five-state model deviates from the simple exponential profile of the one- and two-state models. **c.** The distribution of the average separation (i.e., inverse loaded density) of extruders display a similar level of heterogeneity in the two- and five-state models, which is lacking in the one-state model with immediate rebinding. **d.** The distribution of cohesin extrusion rates, averaged over the chromatin residence time of each extruder, evidences an additional source of heterogeneity not present in one- or two-state models. **e.** Distributions of loop sizes are nonetheless qualitatively similar across the three models.

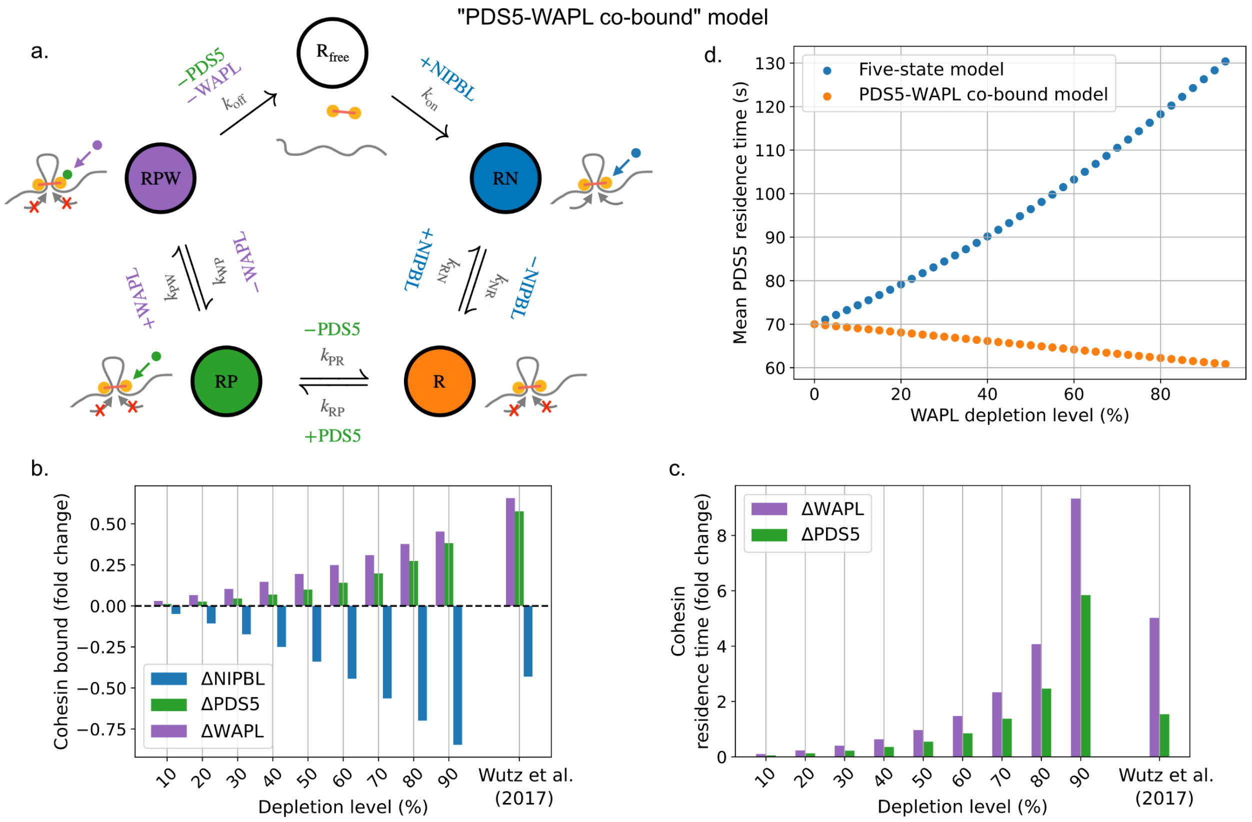


FIG. S5. **A strict co-binding model for cohesin unloading by PDS5 and WAPL.**
**a.** Chemical reaction cycle for the PDS5-WAPL co-bound model. While the reaction cycle is largely similar to that of the five-state model, it differs from the strict regulator exchange considered in the main text by assuming that PDS5 and WAPL can simultaneously bind RAD21, and are jointly required for the unloading of the core complex. **b.** Relative change in cohesin bound fraction with regulator depletion in simulations and experiments (c.f. Fig. 3 of the main text). The simulated effects of PDS5 depletion are now more similar to those of WAPL, while NIPBL depletion has a similar impact to the strict exchange model. **c.** Relative change in cohesin residence time with WAPL or PDS5 depletion. The strictly co-bound model significantly overshoots experimentally-observed increases in cohesin residence time after PDS5 or WAPL RNAi. **d.** Simulated PDS5 residence time as a function of WAPL depletion levels, which suggests the experimental characterization of PDS5 residence time after WAPL depletion (via, e.g., FRAP or single-particle tracking) as a useful metric to assess the validity of the co-bound versus strict exchange models.

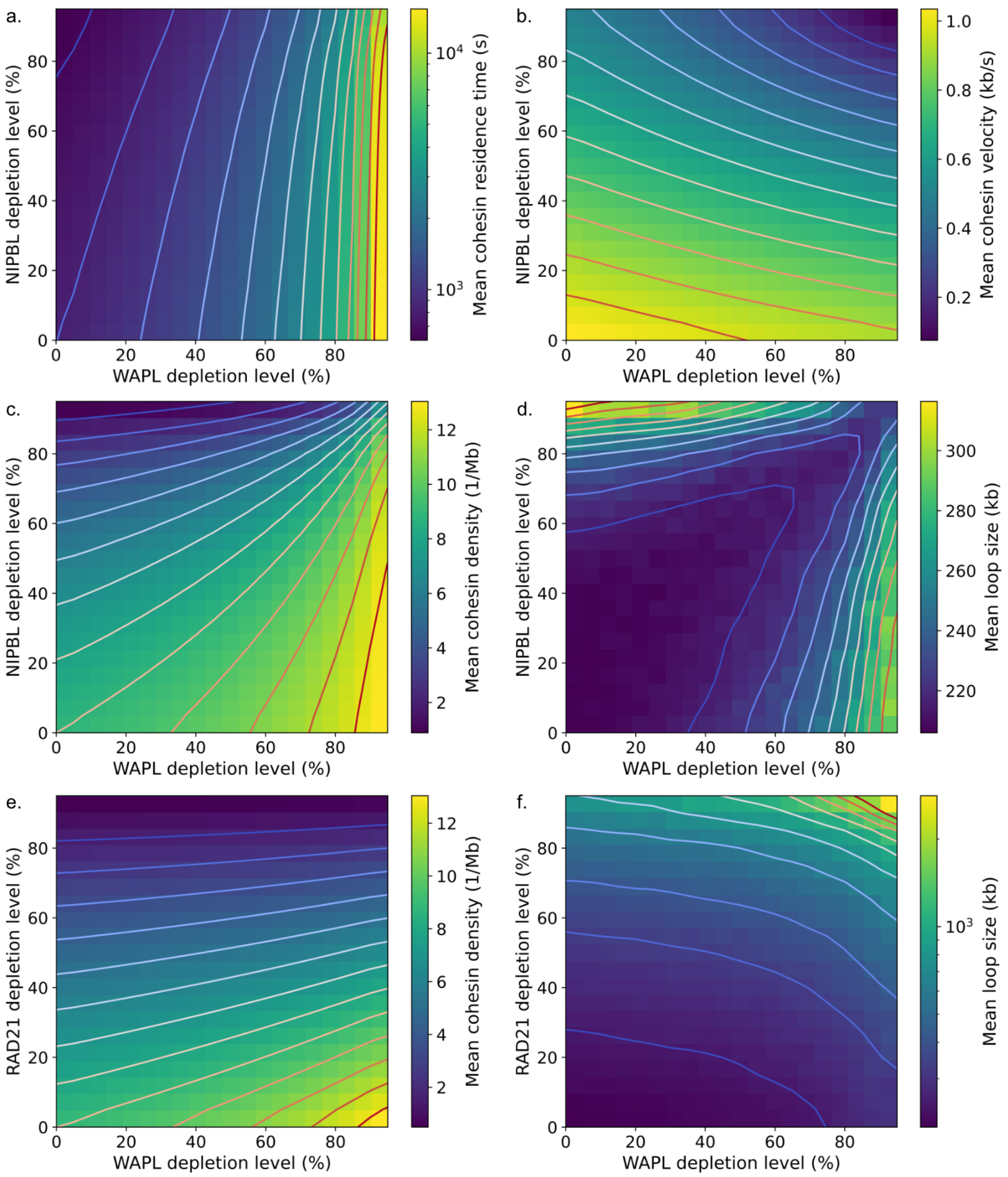
a.
b.
c.
d.
e.
f.
NIPBL depletion level (%)
RAD21 depletion level (%)
WAPL depletion level (%)
Mean cohesin residence time (s)
Mean cohesin velocity (kb/s)
Mean cohesin density (1/Mb)
Mean loop size (kb)

FIG. S6. **Five-state model evidences a compensatory role for NIPBL and WAPL.** **a–d.** Heatmaps indicating changes in various metrics of extrusion activity as a function of NIPBL and WAPL co-depletion levels. Lines show iso-levels of the indicated quantity. NIPBL does not balance the impact of WAPL on residence time, as it only marginally impacts residence time (a). However, NIPBL depletion generally leads to a considerable reduction in the mean translocation rate (b), and can balance the effects of WAPL depletion for both numbers of cohesin per megabase (i.e. cohesin density, c). Thus, NIPBL and WAPL co-depletion generally leads to higher cohesin residence times, but lower extrusion rates — and is thus able to rescue loop sizes when both complexes are down-regulated in similar proportions (d). **e–f.** Same as (c) and (d) for RAD21 and WAPL co-depletion. Unlike NIPBL, RAD21 depletion generally has a limited impact on both residence time and translocation rate (Fig. 3 of the main text), and cannot compensate the effects of WAPL. Thus, while cohesin density can be potentially balanced by simultaneous down-regulation of RAD21 and WAPL (e), loop sizes generally cannot (f).

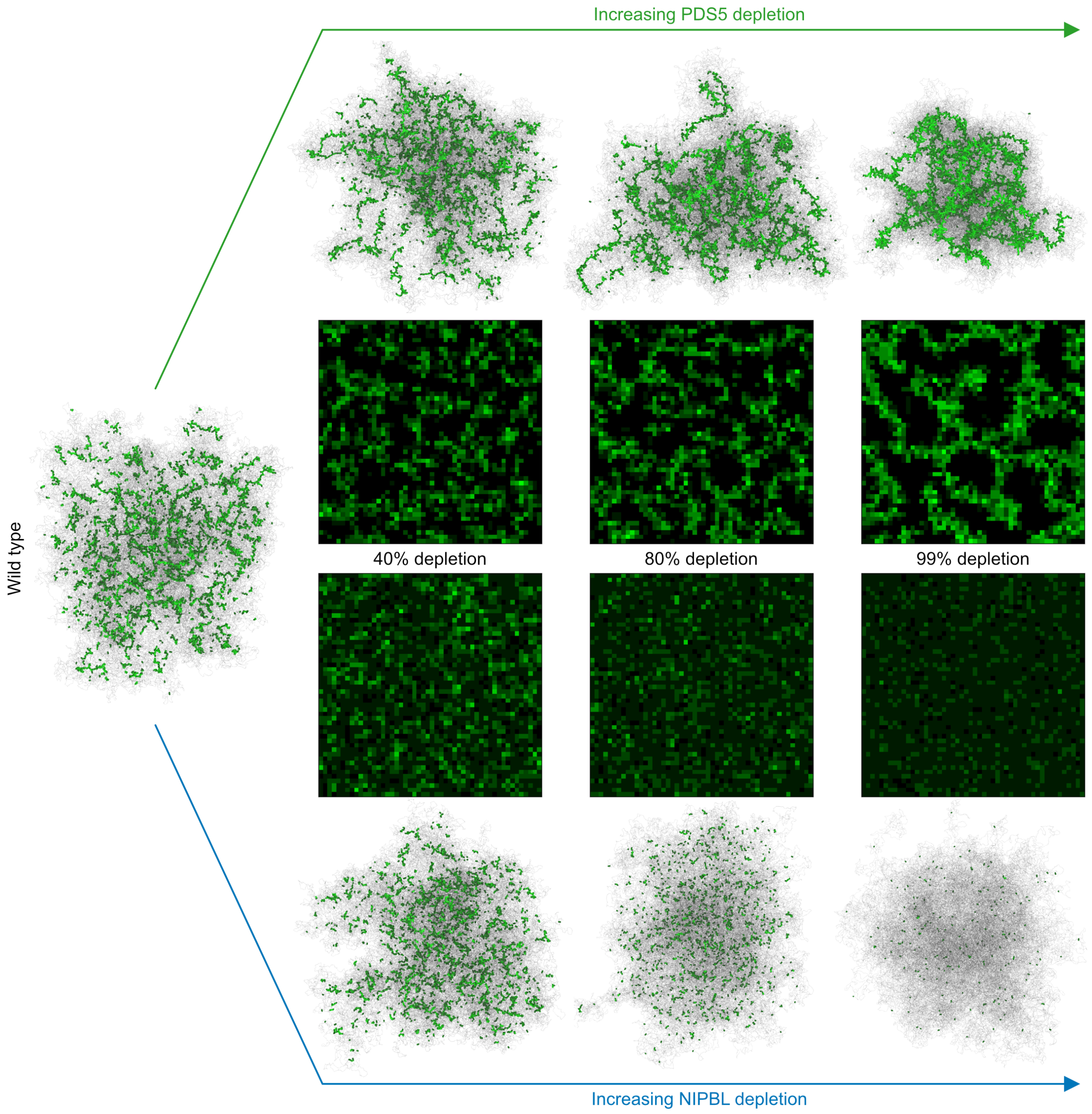


FIG. S7. **PDS5 and NIPBL depletion differentially affect chromosome structure.** Although PDS5 depletion leads to vermicelli phenotypes similar to ΔWAPL (c.f. Fig. 4b of the main text), the reduction in the loaded RAD21 population predicted in the case of NIPBL depletion leads to a gradual disappearance of the chromatin-associated cohesin signal.

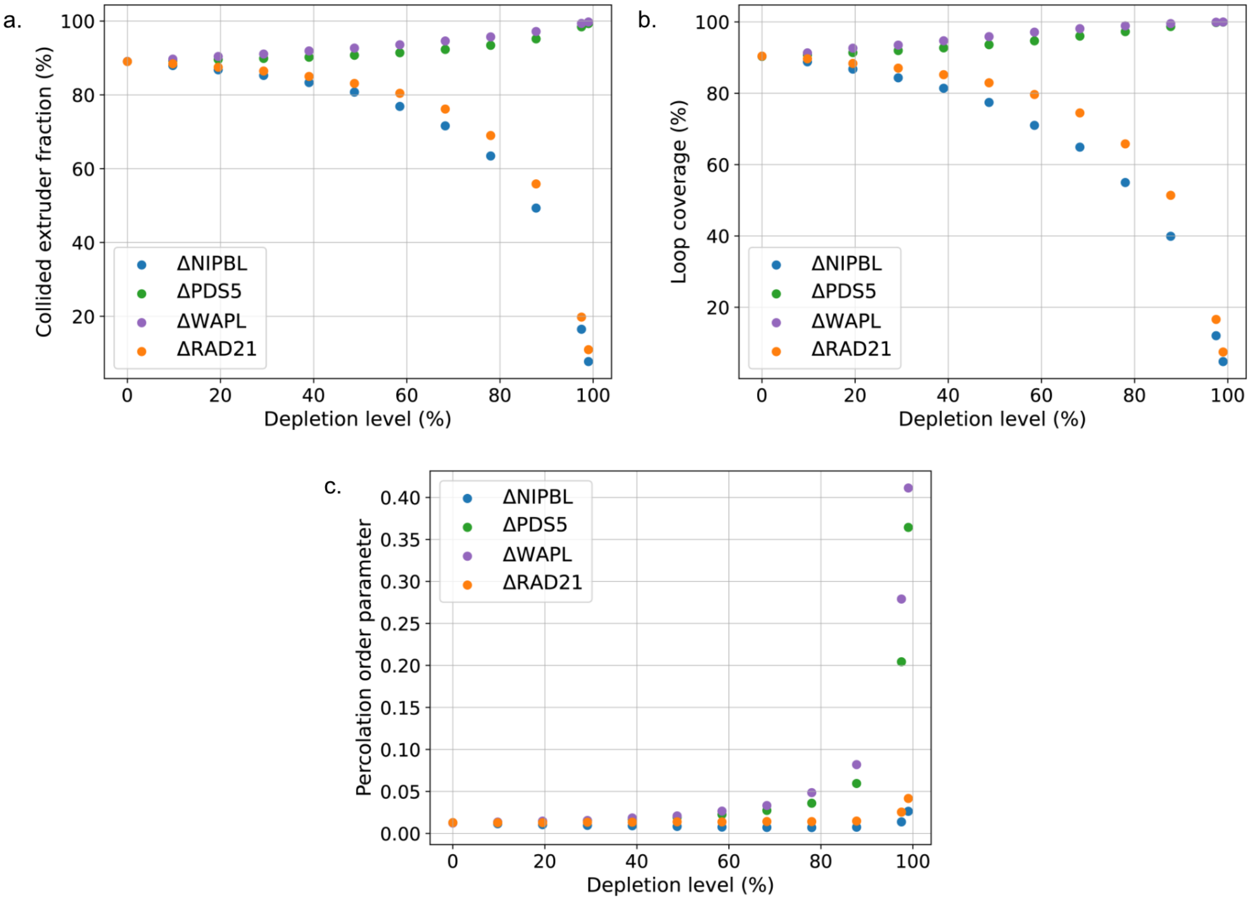


FIG. S8. **Alternative quantification of vermicelli formation.**

**a.** Collided fraction of extruders as a function of depletion for the indicated factor. Collided fraction is calculated as the number of extruder legs directly adjacent to another extruder leg along the 1D lattice, divided by the total number of extruder legs, and are averaged over 5000 lattice conformations obtained across 5 independent simulations. The collided fraction increases with WAPL and PDS5 depletion, approaching 100%, and decreases for NIPBL and RAD21 depletion. **b.** Loop coverage, defined as the fraction of lattice sites that are encompassed by the two legs of any individual extruder. **c.** Cohesin percolation parameter, computed as the size of the largest cluster of collided extruders (as defined in (a)) normalized by the total number of loaded cohesins.

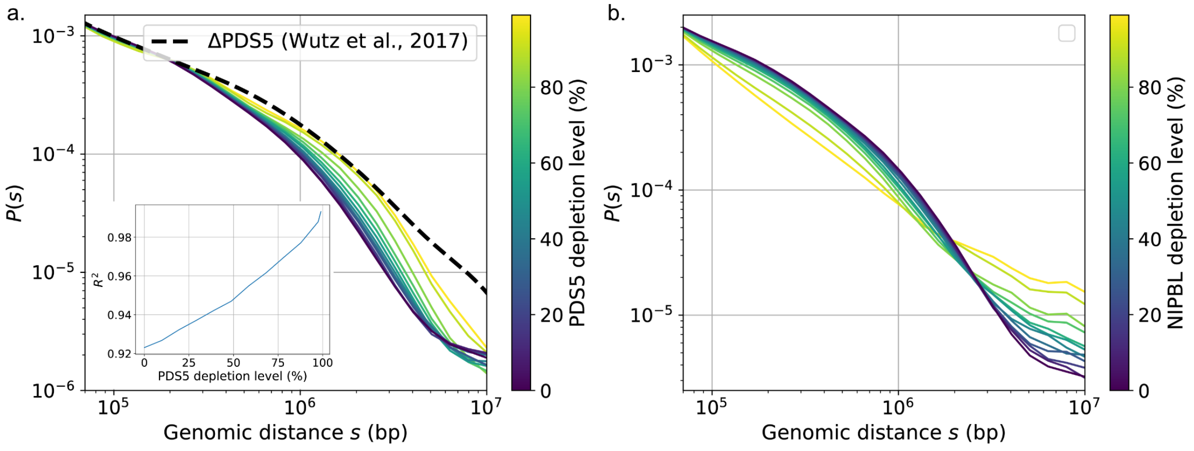


FIG. S9. **Contact frequency versus distance for PDS5 and NIPBL depletion.**
**a.** Simulated PDS5 depletion curves (colored by depletion level) approach experimental data (dashed line) at the highest level of depletion ($\sim$ 99%). **b.** Same as a. for NIPBL depletion. Note that corresponding experimental measurements in NIPBL-depleted HeLa cells are, to our knowledge, currently lacking.

---

[1] G. Wutz, C. Várnai, K. Nagasaka, D. A. Cisneros, R. R. Stocsits, W. Tang, S. Schoenfelder, G. Jessberger, M. Muhar, M. J. Hossain, N. Walther, B. Koch, M. Kueblbeck, J. Ellenberg, J. Zuber, P. Fraser, and J. Peters, EMBO J. **36**, 3573 (2017).

[2] R. Barth, I. F. Davidson, J. van der Torre, M. Taschner, S. Gruber, J.-M. Peters, and C. Dekker, bioRxiv 10.1101/2023.12.21.572892 (2023).

[3] J. Rhodes, D. Mazza, K. Nasmyth, and S. Uphoff, eLife **6**, e30000 (2017).
